# Bayesian Inference of RNA Velocity from Multi-Lineage Single-Cell Data

**DOI:** 10.1101/2022.07.08.499381

**Authors:** Yichen Gu, David Blaauw, Joshua D. Welch

## Abstract

Experimental approaches for measuring single-cell gene expression can observe each cell at only one time point, requiring computational approaches for reconstructing the dynamics of gene expression during cell fate transitions. RNA velocity is a promising computational approach for this problem, but existing inference methods fail to capture key aspects of real data, limiting their utility. To address these limitations, we developed VeloVAE, a Bayesian model for RNA velocity inference. VeloVAE uses variational Bayesian inference to estimate the posterior distribution of latent time, latent cell state, and kinetic rate parameters for each cell. Our approach addresses key limitations of previous methods by inferring a global time and cell state value for each cell; explicitly modeling the emergence of multiple cell types; incorporating prior information such as time point labels; using scalable minibatch optimization; and quantifying parameter uncertainty. We show that VeloVAE significantly outperforms previous approaches in terms of data fit and accuracy of inferred differentiation directions. VeloVAE can also capture qualitative features of expression dynamics neglected by previous methods, including late induction, early repression, transcriptional boosts, and bifurcations. These improvements allow VeloVAE to accurately model gene expression dynamics in complex biological systems, including hematopoiesis, induced pluripotent stem cell reprogramming, neurogenesis, and organogenesis. We find that the latent time automatically inferred using all cells can even outperform pseudotime inferred using manually chosen cell subsets and root cells. We further use the inferred parameters to construct cell type transition graphs and fit branching differential equation models that describe the effects of cell type bifurcations on kinetic rate parameters.

## 1 Introduction

The human body contains many cell types with distinct forms and functions, which arise from progenitor cells in a sequential developmental process. A key question in molecular biology is what regulates this process of cellular development. Therefore, understanding cellular development requires modeling how mRNA expression changes over time. Such models are crucial for numerous areas of biology and medicine, such as neuroscience, cancer research, and regenerative stem-cell therapies.

Since its emergence, the single-cell RNA sequencing (scRNA-seq) technology [32] has been widely used to study cell development. However, scRNA-seq measurement destroys the cell, making it impossible to follow an individual cell longitudinally. Thus, computational approaches are required to assemble these static snapshots into a history of the gene expression changes occuring during a developmental process.

Two main types of computational approaches have been developed for this problem: pseudotime inference and RNA velocity. Pseudotime inference methods use distance from a manually-specified starting cell to rank cells according to degree of development [33, 3]. Many pseudotime methods also aim to infer a tree or graph structure that represents the underlying structure of the developmental process [6, 12, 36, 8, 29]. In contrast, La Manno et al. [19] developed the concept of RNA velocity based on the observation that both unspliced and spliced mRNA molecules appear in sequencing outputs. The relative ratio of spliced and unspliced counts indicates whether the gene was being turned on or turned off at the time the cell was sequenced. La Manno et al. introduced an ODE model to describe the gene expression process, used a steady state assumption to estimate parameters, and implemented the method in a package called velocyto. Later work [4] relaxed the steady-state assumption, allowing all cells to be used in parameter estimation and inferring a latent time value for each cell. Bergen et al. implemented their method in a package called scVelo [4]. We recently extended the dynamical model of scVelo to incorporate chromatin accessibility data, packaged in a tool called MultiVelo [20]. RNA velocity methods have been widely used by biologists to help understand cellular development processes [26, 37, 21].

While they have proven useful for biological discovery in many cases, existing approaches for pseudotime inference and RNA velocity inference have significant limitations. Pseudotime inference requires manually specifying a starting cell, is based purely on pairwise cell similarity, and cannot infer the directions or rates of cell development. RNA velocity addresses some of these limitations, and is in principle able to infer the directions, rates, and origins of developmental processes. However, current RNA velocity methods rely on numerous simplifying assumptions and fail to yield accurate results in many cases [5].

In particular, scVelo suffers from several significant limitations. First, scVelo infers time separately for each gene, which neglects crucial information about the covariance of related genes and often leads to times that are inconsistent across genes. This gene-specific notion of time also makes it hard to compare the switch-off time (time when a cell stops producing new RNA) across genes. The lack of a common time scale, combined with the assumption that induction starts at *t* = 0, also leads to frequent errors in estimating the overall direction of a gene (increasing or decreasing). Genes with a late, short, or missing induction phase are particularly prone to being fit incorrectly by scVelo. Second, scVelo assumes a constant transcription rate *α* within the induction phase for each gene. In a recent review paper, the scVelo developers note that this assumption is often violated in real-world datasets, which leads to a variety of pathological behaviors [5]. Finally, scVelo’s model does not account for cell type bifurcations, which frequently occur in cellular development and can significantly reduce the accuracy of the scVelo model’s predictions.

To address these limitations, we developed VeloVAE, a Bayesian model that uses neural networks to jointly infer the posterior distribution of cell times, cell states, and gene expression rate parameters from scRNA-seq data. Our approach uses a simple, interpretable differential equation model to describe the dynamics of gene expression, but allows the parameters to vary continuously with cell state. The introduction of a cell state variable and a single latent time shared across all genes allows VeloVAE to model qualitative features of expression dynamics neglected by previous methods, including late induction, early repression, transcriptional boosts, and bifurcations. Consequently, VeloVAE can be used as a general tool to reconstruct the orders and rates of gene expression changes across many complex biological systems, including hematopoiesis, induced pluripotent stem cell reprogramming, neurogenesis, and organogenesis.

## 2 Results

### 2.1 VeloVAE Allows Bayesian Inference of Cell Times, Cell States, and Rate Parameters

VeloVAE uses a Bayesian model for RNA velocity inference. We assume that we are given individual scRNA-seq profiles that measure the amounts of spliced (*s*) and unspliced (*u*) transcripts at single moments of a developmental process. Our goal is to use these observations to simultaneously infer the posterior distributions of underlying latent variables that generated the data: cell time (*t*), cell state (**c**), and rate parameters (***θ***) describing the biochemical kinetics of gene expression. Our key modeling assumption is that the observed time-varying (*u*(*t*),*s*(*t*)) levels are related by a system of two ordinary differential equations (Fig. 1a). As with previous RNA velocity approaches, these ODEs capture the simple insight that a gene must first be transcribed as nascent mRNA, then spliced into mature mRNA, and then subsequently degraded (Fig. 1A). However, we make one important change: rather than assuming a single fixed transcription rate parameter for each gene across all cells, we allow each cell to have its own transcription rate *ρ* for each gene. This simple change removes the need for discrete induction and repression phases and models continuous changes in transcription rates, such as transcriptional boosts [5] and cell fate bifurcations. Note also that, unlike scVelo, which places each gene on a separate time scale, the cell time parameter *t* is shared across all genes within a cell.

**Fig. 1.**
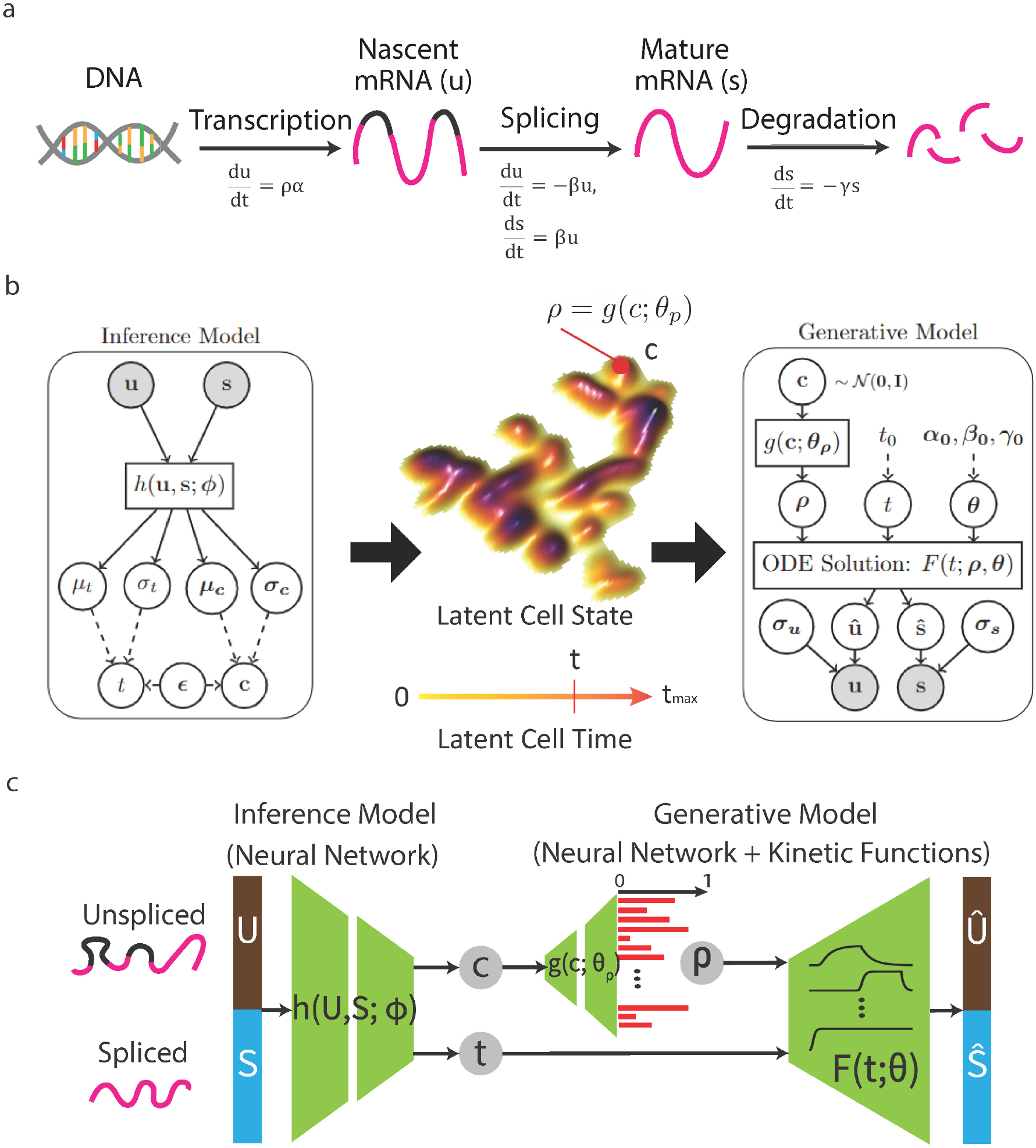
VeloVAE model. **(a)** Differential equation model of transcription used by VeloVAE. Nascent mRNA molecules are transcribed at gene- and cell-specific rate *ρα*. Next, nascent mRNA is spliced into mature mRNA at gene-specific rate *β*. Finally, mature mRNA is degraded at gene-specific rate *γ*. **(b)** Graphical model for latent variable inference and data generative processes. Cell time and state are treated as latent variables, which are inferred using variational Bayes. Note that, unlike previous approaches, latent time (and state) is shared across all genes within each cell. The latent cell state lies on a smooth lowdimensional manifold representing cell development. Each cell state has gene-specific transcription rates *ρ*. RNA count data are generated based on an ODE system with known analytical solution. **(c)** VAE architecture. The encoder neural network learns estimates the posterior distribution of the latent variables, while the decoder learns a mapping from cell state to cell-wise transcriptional rates and simulates the generative process from an ODE system.

The VeloVAE model can be viewed from either an inference or a generative perspective (Fig. 1b). From an inference perspective, if we know (*u*(*t*),*s*(*t*)) values for cells, we can infer something about the (*t*,**c**,***θ***) parameters that generated them (Fig. 1b, left). Each particular location **c** in cell state space has an associated transcription rate *ρ* for each gene; nearby cell state space locations will have similar transcription rates (Fig. 1b, middle). From a generative perspective, if we know the (*t*,**c**,***θ***) parameters for a cell, we can predict the distribution of their (*u,s*) values (Fig. 1b, right). We can also incorporate prior information about the latent variables and rate parameters. If cell capture times are known (e.g., if cells were isolated separately on days 7 and 14), we can use the capture times as an informative prior for cell time.

To fit this statistical model on real data, we use autoencoding variational Bayes, a powerful statistical inference method in which neural networks approximate the posterior distribution of latent variables that may be nonlinearly related to observed data. Intuitively, autoencoding variational Bayes jointly trains the inference and generative models shown in Fig. 1b, so that after training we can infer latent variables given observed data or predict new data given values of the latent variables. VeloVAE implements the inference model of Fig. 1b using a neural network that takes (*u,*s) values as input and outputs the posterior distribution of cell time and cell state parameters (Fig. 1c). VeloVAE implements the generative model of Fig. 1b using a neural network that predicts the gene-specific transcription rates *ρ* for each cell from the cell’s time and state values (Fig. 1c). The (*u,s*) values for each cell can then be predicted using the analytical solution to the ODE, which describes how spliced and unspliced counts vary over time (Fig. 1c). We previously described the underlying theory and motivation of this approach in a machine learning conference paper [10]. Intuitively, VeloVAE is like an autoencoder whose decoder network has been replaced with the solution to a differential equation. Importantly, we also take advantage of the little-known fact that autoencoding variational Bayes allows inference of posterior distributions for parameters in the generative model (decoder network). This allows us to infer distributional estimates of the ODE rate parameters as well.

We train the model by maximizing the evidence lower bound of the marginal likelihood using mini-batch stochastic gradient descent [15]. Crucially, cells are loaded in batches during training, which means that not every cell must be used in each training iteration, dramatically decreasing both the time and memory requirements.

### 2.2 VeloVAE Significantly Improves Model Fit and Latent Time Accuracy

We evaluated our method on a variety of real scRNA-seq datasets of different sizes and complexity [2, 25, 28, 31, 18, 6] and compared our results with scVelo [4], the state-of-the-art method for RNA velocity inference. We evaluated VeloVAE and scVelo in terms of both how well the models fit the data and how accurately they recover latent time. To quantify model fit, we calculated mean squared error (MSE), mean absolute error (MAE), and log likelihood (LL) as our metrics (Fig. 2a, S1). For VeloVAE, we calculated these metrics on both the training dataset and a held-out test set not used during training. Note that scVelo cannot perform out-of-sample prediction, so we were not able to evaluate it on a held-out test set. Our results show that VeloVAE achieves much better data fit than scVelo for all datasets in the evaluation. Furthermore, this better performance does not come from overfitting the training dataset - VeloVAE shows similarly good performance on the held-out datasets.

**Fig. 2.**
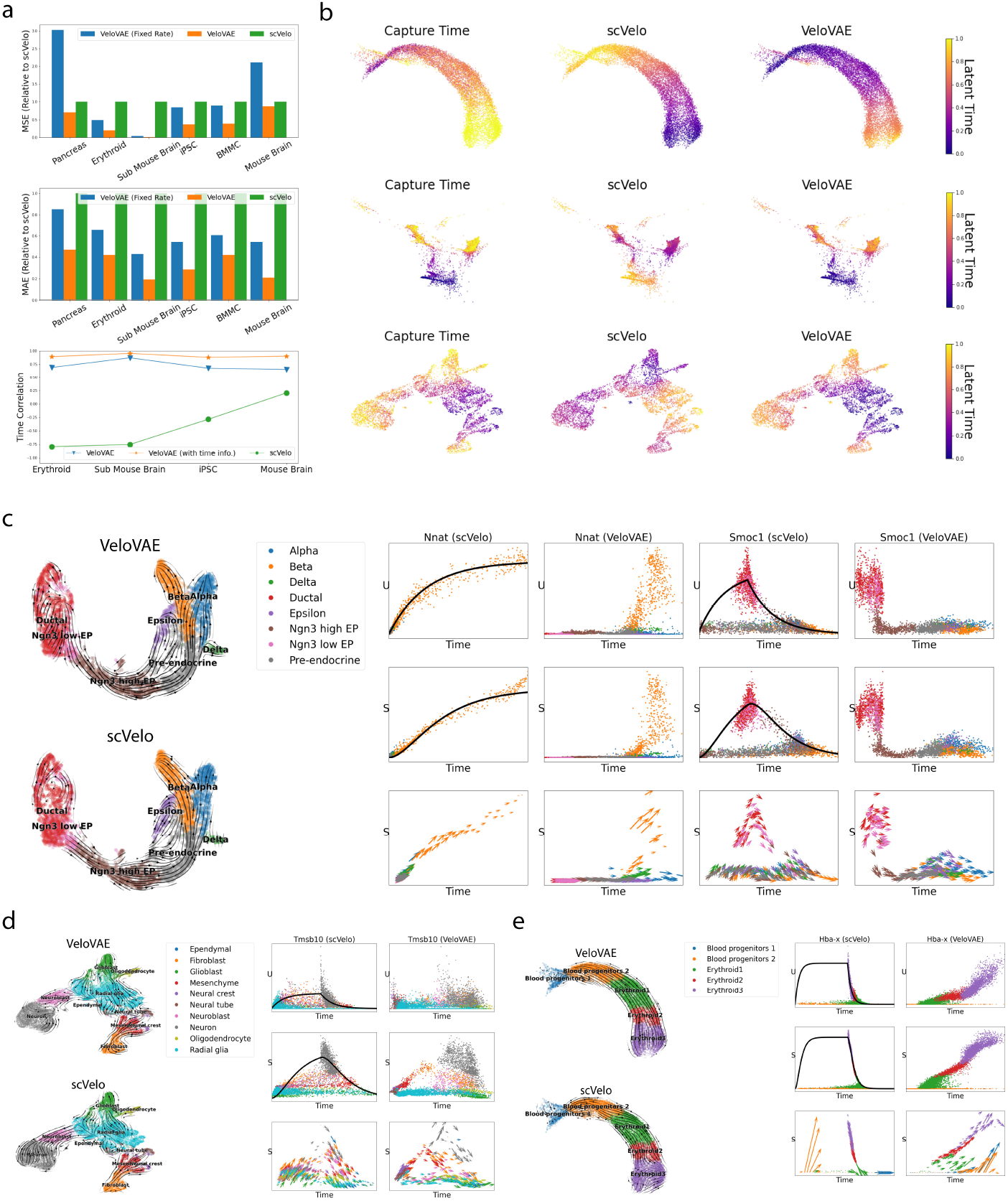
VeloVAE Significantly Improves Model Fit and Latent Time Accuracy and Models Complex Gene Expression Dynamics. **(a)** Quantitative performance comparison. We report the mean squared error (MSE) and mean absolute error (MAE) between the observed and predicted (u, s) counts, as well as correlation between inferred latent time and true capture time (when available). MSE and MAE are reported relative to scVelo. **(b)** UMAP plots colored by true capture time and inferred latent time for erythroid, iPSC, and subsampled mouse brain datasets. **(c)-(e)** UMAP plots and examples of individual genes fit by scVelo and VeloVAE for pancreas **(c)**, subsampled mouse brain **(d)**, and erythroid **(e)** datasets. UMAP plots are colored by published cluster assignments with inferred velocity streams overlaid. Gene fits are shown for both *u* and *s* values, with inferred latent time on the x-axis. Fitted values from scVelo are shown as lines, with observed data values shown as points. Only fitted values are shown for VeloVAE, because the VeloVAE fit is a point cloud (rather than a line) that would completely cover the observed data. Points are colored by published cluster assignments, with the same colors as in the corresponding UMAP plots. Note that the x-axis values are different for scVelo and VeloVAE because they infer different latent times. The arrow length and direction indicate the velocity inferred for each cell.

To evaluate the accuracy of latent time inference, we used scRNA-seq datasets with cells sampled from multiple time points. These time points usually have rather coarse granularity (e.g., day 7 and day 14), and cells captured at the same time may span a wide range of developmental stages. Nevertheless, the inferred cell times should at least be correlated with the capture times. Thus, we computed the Spearman correlation between the cell times inferred by each method and the capture times (Fig. 2a). VeloVAE consistently and significantly outperforms scVelo in terms of latent time accuracy, with scVelo often inferring latent time that is anticorrelated with real time (Fig. 2a,b). VeloVAE achieves a significant improvement in latent time accuracy even without using the time labels, though when the time labels are used as an informative prior, the time correlation improves further (Fig. 2a, bottom). Although scVelo infers latent time separately for each gene, the tool provides a post-hoc procedure for estimating a single global time for each cell. Using this global time for comparison with our methods casts scVelo in the best possible light because the global time is more robust than the gene-specific latent times. The low time correlation from scVelo may be partly explained by inconsistency among the different notions of time fitted for each gene. To investigate this further, we computed the average time correlation between scVelo’s gene-specific and global latent time. As Figure S2 shows, the correlation between scVelo’s global latent time and the latent time for each gene is indeed quite low; the latent time values for many genes are even anticorrelated with global latent time.

We further assessed the relative importance of using a shared latent time and using a cell state variable. To do this, we fit a version of VeloVAE that has only a shared latent time but no cell state variable. The performance was better than scVelo in many cases due to the shared latent time, but generally significantly worse than the model with the cell state variable (Fig. 2a). This indicates that including a transcription rate that varies with cell state is crucial for the best performance.

### 2.3 VeloVAE Better Captures Qualitative Properties of Complex Gene Expression Dynamics

We designed the VeloVAE model to relax several of the restrictive assumptions of previous RNA velocity approaches. Thus, we expect that the model should show increased expressiveness, allowing it to capture qualitative properties of gene expression changes that previous approaches cannot. To assess this, we fit both VeloVAE and scVelo on three representative datasets–mouse pancreas, human blood, and mouse brain–and inspected the resulting model fits. We found three types of qualitative behaviors that scVelo and previous approaches cannot accurately model, while VeloVAE can.

#### Late Induction and Early Repression

We observed that genes with a late, short, or missing induction phase are particularly prone to being fit incorrectly by scVelo. The lack of a common time scale, combined with the assumption that induction starts at *t* = 0, also leads to frequent errors in estimating the overall direction of a gene (increasing or decreasing). A dataset from the mouse pancreas [2] illustrates this behavior.

VeloVAE and scVelo both yield latent time values and stream plots for the pancreas dataset that are coherent with prior knowledge (Fig. 2c). However, inspecting the fits for individual genes shows that scVelo often rearranges the local time for each gene to try to force the genes to have an induction phase starting at t = 0. Figure 2c shows two sample genes, *Nnat* and *Smoc1*. The *Nnat* gene is not turned on until the pre-endocrine cells appear, whereas *Smoc1* is immediately switched off at the beginning of the differentiation process. We can see that scVelo fails to detect late induction in *Nnat* and assigns the latent time to zero for almost all cell types except for beta cells. For *Smoc1*, scVelo rearranges the order of progenitor and descendent cell types in an effort to force the gene to have a induction phase. In contrast, VeloVAE is able to correctly infer the late induction pattern for *Nnat* and the early repression pattern for *Smoc1*.

#### Cell Type Bifurcations

Many cell differentiation processes produce multiple descendant cell types from a single progenitor type, but previous RNA velocity approaches model only a single cell type. By including a cell-specific latent state, VeloVAE can model the continuous emergence of multiple cell types from a single progenitor type. For example, in the pancreas dataset, the *Nnat* gene is upregulated as cells differentiate towared the beta cell fate, but not in any other cell type (Fig. 2c).

As another example, we fit both scVelo and VeloVAE on a developing mouse brain atlas [18]. For clarity, we subsampled the dataset to include only cell types arising from the neural tube (see below for an analysis of the full dataset). In this system, neural tube cells develop into radial glia. Some of the radial glia cells differentiate into neuronal cell types, while the others give rise to the glial lineage, including glioblasts, oligodendrocytes, astrocytes, and ependymal cells. In short, this is a complex system with many distinct lineages emerging. Nevertheless, VeloVAE can accurately model the complex, multi-lineage dynamics of genes in this system. For example, VeloVAE accurately models the behavior of the *Tmsb10* gene (Fig. 2d), which is turned on in the fibroblast and neuronal cells at different times and remains off in the non-neuronal cells differentiated from radial glia. In contrast, scVelo rearranges the latent time values of the cells in a vain attempt to fit the *Tmsb10* gene into a single induction and repression cycle.

#### Transcriptional Boosts

In addition to bifurcation, cell differentiation can involve gradual or abrupt increases in transcription rate within a cell type. Several recent papers reported that erythroid cell differentiation involves abrupt “transcriptional boosts” [5,1]. Previous RNA velocity approaches assumed a constant transcription rate, and thus could not model this behavior. Indeed, when we fit scVelo on an erythroid dataset, we find that scVelo infers latent time that is strongly anticorrelated with true time (Fig. 2b,e). Similarly, scVelo predicts that the *Hba-x* gene, which shows a transcriptional boost, is repressed rather than induced. In contrast, VeloVAE is able to model the transcriptional boost in *Hba-x* levels (Fig. 2e) because the transcription rate *ρ* is cell-specific.

### 2.4 Cell Type Transition Graph Inference and Branching Differential Equation Model

Understanding cell differentiation processes requires discovering which progenitor types give rise to which differentiated types. In addition, we want to know how the expression of key fate determining genes changes during cell type bifurcation. We reasoned that the VeloVAE results provide several opportunities to investigate these questions.

Although VeloVAE accurately models bifurcations using a continuous cell state variable, the resulting parameters are not readily interpretable in terms of discrete cell types. Thus, we developed a model extension that aids in interpreting how the kinetic parameters of gene expression change across cell types. We extended the simple differential equation model shown in Fig. 1 to a system of equations that we refer to as a branching ODE model (Fig. 3a). Instead of a cell-specific transcription rate and fixed splicing and degradation rates, the branching ODE model assigns each cell type a unique ODE with cell-type-specific transcription, splicing, and degradation rates and an initial condition determined by the progenitor cell type (Methods). The model relies on a directed graph relationship among discrete cell types. The graph can be inferred directly from the data, as we did here; if some aspects of the cell type transition graph are known, these can also be manually specified. We constrain the branching ODE model so that each cell type emerges at a specific time and the initial conditions of each cell type match the ODE prediction of its parent cell type at the time the child cell type emerges. This provides a qualitative view of the change of kinetic rates during cell development. We use the branching ODE model to replace the decoder of the VeloVAE, giving an alternative, more interpretable generative model.

**Fig. 3.**
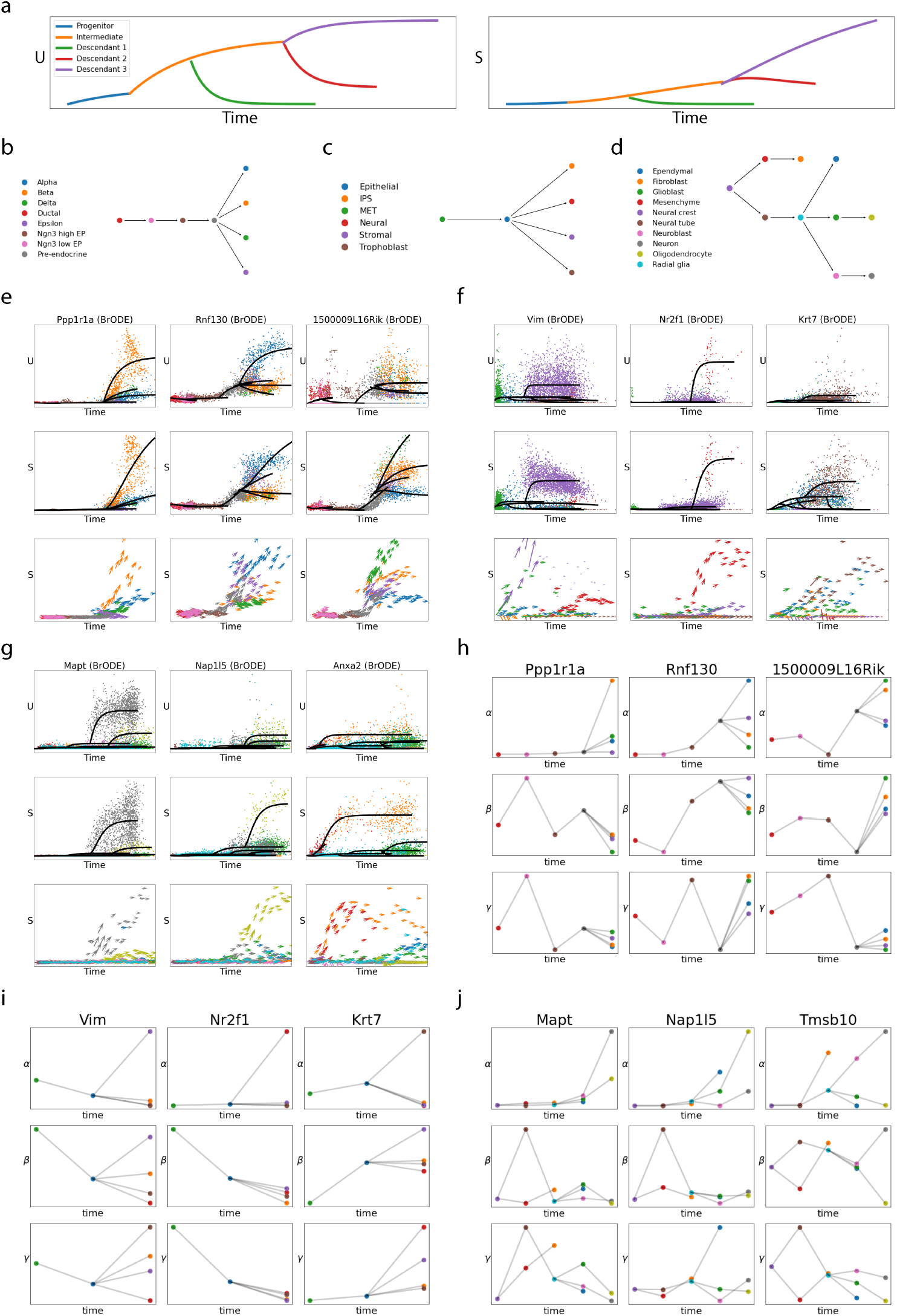
Cell Type Transition Graph Inference and Branching Differential Equation Model. **(a)** Schematic of branching ODE solutions. The example shows the output prediction of *u* and *s* versus time for a process with a single progenitor, an intermediate and three distinct descendant cell types. **(b)-(d)** Inferred cell type transition graphs from pancreas **(b)**, iPSC **(c)** and subsampled mouse brain **(d)** datasets. In each graph, a vertex represents a cell type and a directed edge points from a progenitor cell type to its immediate descendant(s). **(e)-(g)** Examples of individual genes fit by the branching ODE model for pancreas **(e)**, iPSC **(f)** and subsampled mouse brain **(g)** datasets. Each column represents a gene and plots the unspliced count, spliced count and RNA velocity versus time from top to bottom. The branching ODE fits are shown as solid lines. **(h)-(j)** cell-type-specific rate parameters inferred by branching ODE model for pancreas **(h)**, iPSC **(i)** and subsampled mouse brain **(j)** datasets. Each column represents a gene and plots the transcription (*α*), splicing (*β*) and degradation (*γ*) rates from top to bottom. Each point represents a cell type, the types are arranged in chronological order, and progenitor-descendant relationships are indicated with lines.

Having established the accuracy of the results from VeloVAE, we used these results to infer cell type transition graphs. To infer the transition probability between cell types A and B, we used the cell times to simply count how often cells of type A and B occur in two immediately adjacent short time intervals (see Methods for details). We show three examples of these graphs (Fig. 3b-d), inferred from the pancreas, iPS reprogramming, and mouse brain datasets from Fig. 2. The inferred cell type transition graphs match closely with biological expectations for these systems, even for the complex mouse brain dataset.

To train the model, we first obtain the time and state assignments for each cell by training VeloVAE. Next, we infer a transition graph describing the progenitor-descendant relations among cell types (Methods). Finally, we fix the encoder of VeloVAE (so that latent time and cell state estimates are fixed) and estimate the parameters of the branching ODE model. We perform this parameter estimation by maximizing the Gaussian likelihood of all genes under the branching ODE model, which is equivalent to minimizing the Mahalanobis distance between model fit and observed data.

We trained branching ODEs on pancreas, iPSC and subsampled mouse brain datasets. The cell type transition graphs are all consistent with prior knowledge (Fig. 3b-d). For example, our computational method successfully finds the expected cell differentiation path in pancreatic development, starting from ductal cells and branching into *α, β, δ* and *ϵ* cells (Fig. 3b). Importantly, we achieved these accurate results from simply training VeloVAE using default parameters without any parameter tuning; we used the same parameters for all three of these datasets.

In addition, the branching ODE is able to infer different and asynchronous gene expression kinetics in multiple branches and successfully captures different rates in different branches in these datasets. For example, *Ppp1r1a* has almost all transcription activity in β cells (Fig. 3e), which was verified by previous studies [14, 7]. The *Rnf130* and *1500009L16Rik* genes similarly show significant branching trends that are accurately modeled by the branching ODE (Fig. 3e). The branching ODE model also accurately fits genes that show differential kinetics among lineages that emerge during induced pluripotent stem cell reprogramming. For example, *Vim* is strongly upregulated as epithelial-like cells transition to stromal-like cells (Fig. 3f). The *Nrf21* gene is strongly and specifically upregulated in neural-like cells (Fig. 3f). The *Krt7* gene is upregulated in epithelial-like and trophoblast-like cells with differing expression levels (Fig. 3f).

Another example is the *Mapt* gene from the mouse brain dataset (Fig. 3g). The gene is upregulated strongly in neurons and subsequently transcribed at much lower levels in oligodendrocytes, which coheres with previous studies [17]. The *Napl15* gene shows the opposite trend, with high transcription in oligodendrocytes and low but detectable transcription in glioblasts and neurons (Fig. 3g). As another example, the *Anxa2* gene is most highly transcribed in the early transition from mesenchyme to fibroblast, with some later transcription in glioblasts and ependymal cells (Fig. 3g). The cell-type-specific rate parameters inferred by fitting the branching ODE describe the differences in transcription, splicing, and degradation that cells undergo as they differentiate (Fig. 3h-j).

### 2.5 VeloVAE Accurately Models Human Hematopoiesis

Previous papers have noted that hematopoiesis is a particularly difficult system for existing RNA velocity methods [5]. Latent time and velocity inferences often seem to point in the opposite direction of the known blood cell differentiation hierarchy. Two aspects in particular likely make this system challenging. First, many distinct cell types emerge simultaneously from the hematopoietic stem cell (HSC). Recent studies suggest that hematopoietic progenitors are more like a continuum of primed states than a set of discrete states neatly organized in a tree structure [35]. Second, blood cells are produced exceptionally rapidly compared with other cell types; a recent study estimated that about 2 million new red blood cells per second enter the bloodstream [13]. Thus, blood cells may use special gene regulatory mechanisms such as “transcriptional boosts” (see discussion above) and other time-varying rate parameters.

To investigate whether VeloVAE can resolve these difficulties, we analyzed a recent human bone marrow dataset [31]. Our stream plot shows that VeloVAE correctly identifies HSCs as the start of differentiation and predicts that they differentiate into megakaryocytes, platelets, dendritic cells, monocytes, and B-cells (Figure 4a). In contrast, the scVelo stream plot predicts incorrect differentiation vectors that point backward for the B-cell, dendritic cell, platelet, and megakaryocyte lineages (Fig. 4b). Inspecting the fits of individual genes revealed that scVelo fails to capture the complexity of multiple lineages simultaneously emerging, as well as time-varying transcription rates (Fig. 4c). For example, the *TCF4* gene is upregulated strongly in pDCs and moderately in B cell progenitors, but scVelo incorrectly infers that the gene is globally downregulated, while VeloVAE correctly models the expression dynamics (Fig. 4c). This reversed trend predicted by scVelo matches the reversed scVelo streamplot observed for pDCs and B cell progenitors, suggesting that multiple genes are likely fit in a similarly incorrect manner. As another example, the *FHIT* gene shows complex, multi-lineage kinetics with distinct expression levels in T- and B-cell lineages. VeloVAE can accurately model this behavior, while scVelo tries to collapse the expression levels into a single global trend by rearranging cell times (Fig. 4c). Latent time from both methods is similar for many cells, but VeloVAE is clearly more accurate for the rare differentiated populations, such as dendritic cells, platelets, and plasmablasts (Fig. 4d).

**Fig. 4.**
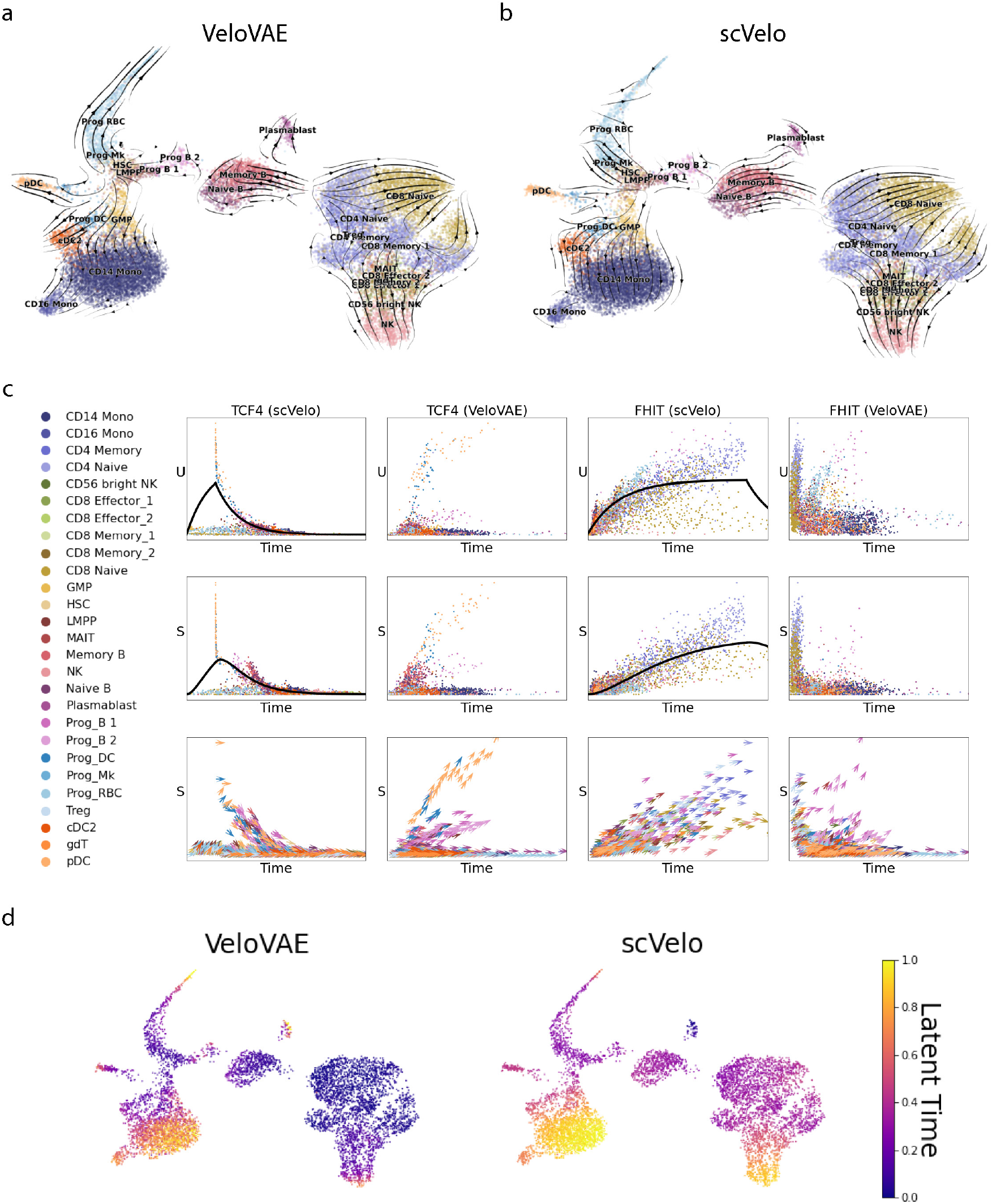
VeloVAE Models Complex Kinetics in Bone Marrow Cells. UMAP plots colored by published cell types overlaid with velocity inferred by **(a)** VeloVAE or **(b)** scVelo. **(c)** Examples of individual genes fit by scVelo and VeloVAE. Gene fits are shown for both *u* and *s* values, with inferred latent time on the x-axis. Fitted values from scVelo are shown as lines, with observed data values shown as points. Only fitted values are shown for VeloVAE, because the VeloVAE fit is a point cloud (rather than a line) that would completely cover the observed data. Points are colored by published cluster assignments, with the same colors as in the corresponding UMAP plots. Note that the x-axis values are different for scVelo and VeloVAE because they infer different latent times. The arrow length and direction indicate the velocity inferred for each cell. **(d)** UMAP plots colored by latent time inferred by VeloVAE and scVelo.

### 2.6 VeloVAE Accurately Models Cell Differentiation Across the Whole Mouse Brain

Single-cell technologies now enable large-scale measurement of differentiating cells across whole organs or organisms, but modeling the emergence of cell fates at that scale remains extremely challenging with existing computational approaches. For example, La Manno et al. recently published an atlas of the entire developing mouse brain, but did not perform any RNA velocity analysis [18]. The dataset contains 292,495 cells of 22 major types and 798 subtypes, extracted between embryonic days 7 and 18. Starting from late gastrulation, the neuronal lineage develops from neural tube and neural crest cells. Neural crest cells differentiate into fibroblasts, while neural tube cells differentiate into radial glia cells, which then branch into both neuronal and glial cell types.

As Figure 5a and b show, VeloVAE captured the cellular dynamics in all major lineages, while the dynamical model from scVelo failed to capture the correct dynamics in the neuronal lineage. This is also manifested in the gene expression dynamics (Fig. 5c). Furthermore, the latent time from VeloVAE is qualitatively close to the cell capture times, reflecting the true time order of the cells (Fig. 5d).

**Fig. 5.**
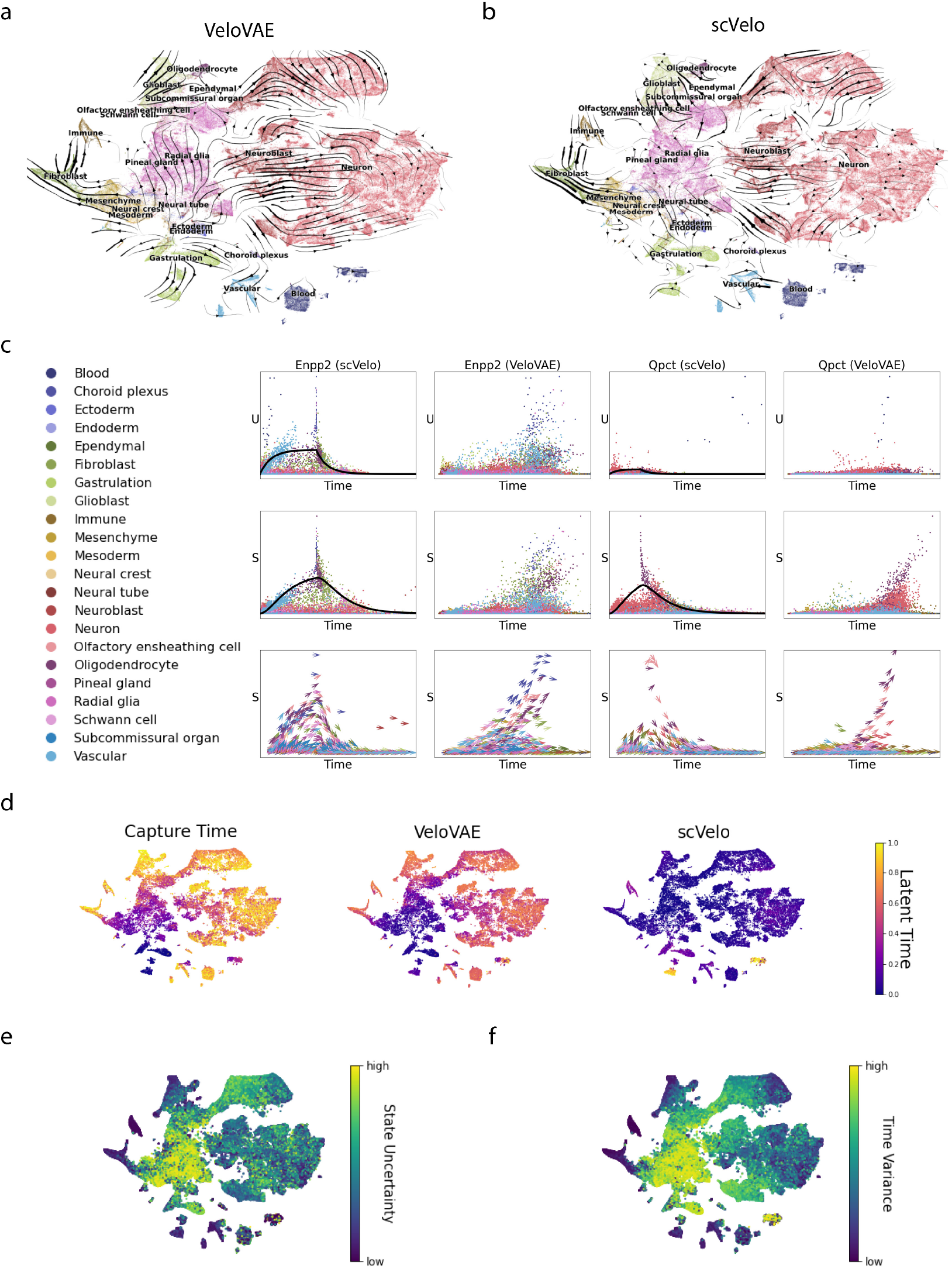
VeloVAE Resolves Cellular Dynamics of Multiple Lineages Across the Entire Developing Mouse Brain. t-SNE plots colored by published cell types overlaid with velocity inferred by **(a)** VeloVAE or **(b)** scVelo. **(c)** Examples of individual genes fit by scVelo and VeloVAE. Gene fits are shown for both *u* and *s* values, with inferred latent time on the x-axis. Fitted values from scVelo are shown as lines, with observed data values shown as points. Only fitted values are shown for VeloVAE, because the VeloVAE fit is a point cloud (rather than a line) that would completely cover the observed data. Points are colored by published cluster assignments, with the same colors as in the corresponding UMAP plots. Note that the x-axis values are different for scVelo and VeloVAE because they infer different latent times. The arrow length and direction indicate the velocity inferred for each cell. **(d)** t-SNE plots colored by true capture time and latent time inferred by VeloVAE and scVelo. **(e)-(f)** t-SNE plots colored by the cell state uncertainty **(e)** and cell time variance **(f)** from VeloVAE.

In addition to the velocity and latent time inferred by VeloVAE, we also analyzed the inferred cell states. Recall that the latent cell state inferred by VeloVAE is a low-dimensional embedding of mRNA counts of each cell. To visualize these cell states, we generated 3D plots where the x-y plane is the 2D UMAP [23] coordinates of cell states and z axis corresponds to the cell time (Supplementary Figure S3f). As a comparison, we generated a 3D quiver plot whose x-y plane is 2D t-SNE [22] coordinates provided by the authors and z axis is still the cell time (Supplementary Figure S4f). In this way, we can visualize the whole differentiation process. In addition, our variational posterior also contains the variance of the cell state. We define the cell state uncertainty as the multi-variate coefficient of variation (CV) [34] of the cell state and visualize it in a t-SNE plot (Fig. 5e, Supplementary Figure S5f). Similarly, we obtained time variance for each cell (Fig. 5f, Supplementary Figure S6f).

From these plots, we noticed an interesting phenomenon: multi-potent progenitor cells often have very high cell state uncertainty. For example, radial glia cells in mouse brain development can differentiate into both neuronal and non-neuronal types, so they have high cell state uncertainty. In contrast, the more differentiated cell types show relatively lower cell state uncertainty. This suggests that the cell state distributions learned by VeloVAE can capture biologically meaningful uncertainty in the states of cells undergoing fate decisions.

### 2.7 VeloVAE Accurately Models Organ Development in Whole Mouse Embryos

The challenges of studying cell differentiation with single-cell data become particularly acute at the scale of an entire organism. For example, Cao et al. performed single-cell RNA-seq on 61 entire mouse embryos sampled from E9.5-E13.5, the developmental period when mouse organogenesis occurs [6]. The dataset contains 1,380,824 cells after preprocessing, which fall into 38 major cell types and can divided into 10 major lineages. Pseudotime analysis and RNA velocity analysis have been performed on this dataset, but both analyses were highly manual processes that required separate curation of dozens of cell subsets [6, 27].

In principle, the continuous cell state variable of VeloVAE is sufficiently expressive to provide a single model of the differentiation potential for an entire organism. To investigate this, we trained VeloVAE on the mouse organogenesis dataset. Because of the large size and cellular diversity of this dataset, we used a larger batch size of 2048 and increased the dimension of **c** from 5 to 10. Our results show that VeloVAE discovered a meaningful latent cell state space representing the whole development process (Fig. 6a). To examine the results in more detail, we ran UMAP individually on the same 10 broad cell lineages that were identified in the initial paper. Importantly, we performed this subset analysis purely for visualization purposes–the cell time, cell state, and kinetic rate parameters were estimated only once using all cells jointly (Supplementary Figure S7–S12).

**Fig. 6.**
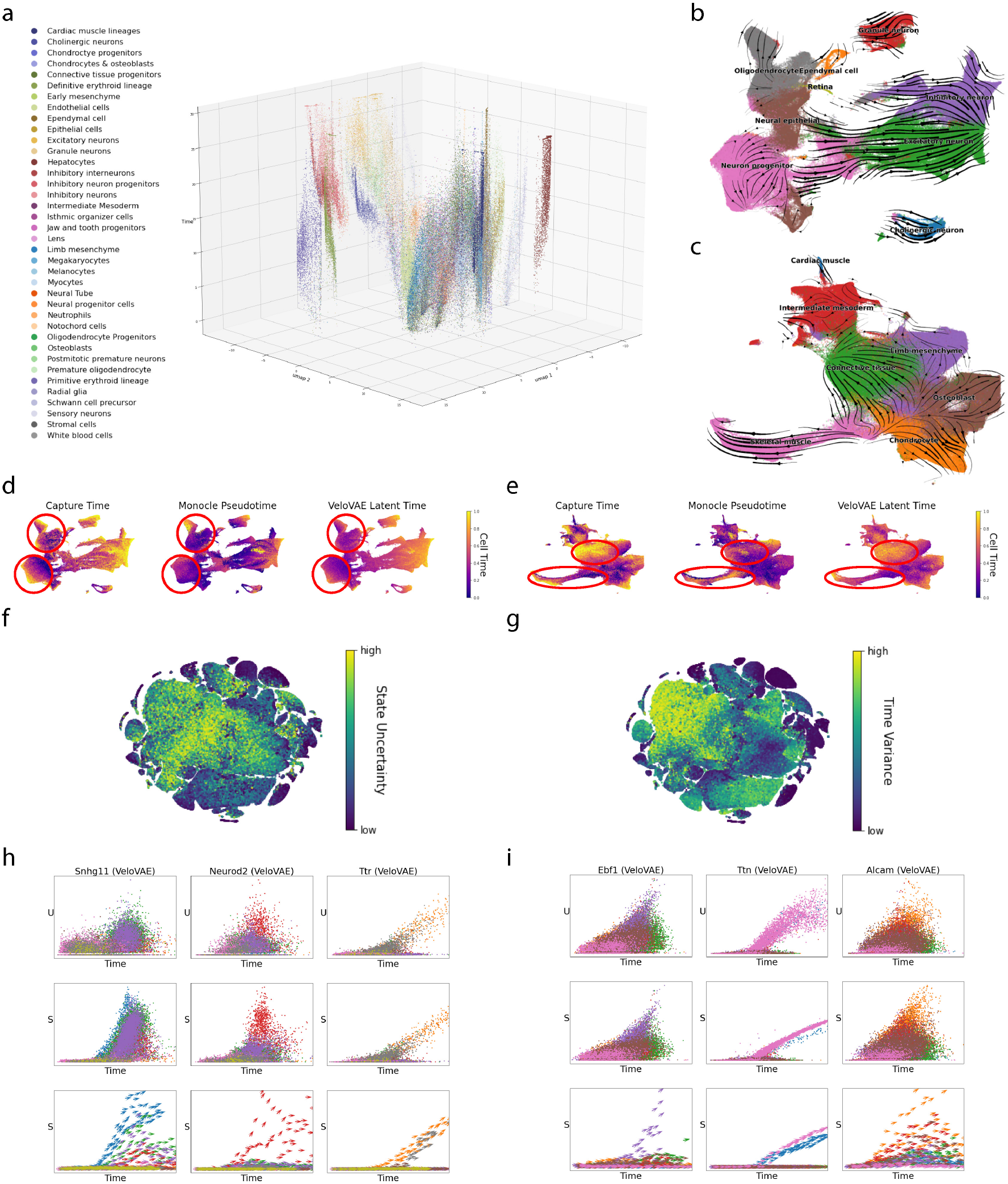
Whole mouse embryo analysis. **(a)** 3D representation of entire mouse embryo dataset, colored by broad cell class. The vertical axis is latent time value inferred by VeloVAE. The horizontal axes are a 2D UMAP projection of the cell states inferred by VeloVAE. **(b)-(c)** UMAP plots colored by published cell types overlaid with velocity inferred by VeloVAE for neuronal **(b)** and mesenchymal **(c)** trajectories. The UMAP coordinates are computed using only cells from each trajectory, but the velocity estimates are from a single VeloVAE model fit on all cells. **(d)-(e)** UMAP plots colored by capture time, published pseudotime, and VeloVAE latent time values for the neuronal **(d)** and mensenchymal **(e)** trajectories. Note that latent times are from a single VeloVAE model fit on all cells. **(f)-(g)** t-SNE plots colored by **(f)** cell state uncertainty and **(g)** time variance. **(h)-(i)** VeloVAE fits for selected genes from the mesenchymal **(h)** and neuronal **(i)** trajectories.

These visualizations indicate that latent time and velocity estimates are highly consistent with cell capture times and biological prior knowledge (Fig. 6b-e, Supplementary Figure S7,S8). Remarkably, the latent time estimates are even more accurate than the authors’ reported pseudotime values in several cases. For example, the VeloVAE latent time estimates for the two lineages with the most cells, mesenchyme and neural tube, both show a higher correlation with capture time than the pseudotime estimates reported by Cao et al [6]. In particular, the pseudotime estimates for the oligodendrocyte and neural progenitor cells are essentially uncorrelated with capture time, whereas the VeloVAE results proceed in the direction of increasing embryonic stage (Fig. 6d). Similarly, the pseudotime estimates for skeletal muscle cells are actually anticorrelated with cell capture times, and the pseudotimes of developing connective tissue cells substantially underestimate their developmental progress (Fig. 6e).

These results are remarkable because the pseudotime values were generated by the developers of Monocle3–one of the most popular pseudotime methods–on their own data using their own tool in a highly manual process. Cao et al. used expert knowledge to group cells into “subtrajectories” and choose root cells for each. In contrast, we obtained the VeloVAE results by simply running the model jointly on all cells with no manual curation. The accuracy of the VeloVAE latent time estimates underscores the power of this approach for studying cell differentiation in large, complex, multilineage single-cell datasets. This comparison also highlights the inherent difficulty of pseudotime analysis in such complex datasets, particularly when the continuous nature of differentiation and the presence of multiple “root cell” populations make it very challenging to divide the cells into discrete subtrajectories.

In comparing latent time and pseudotime values, we realized that the latent time values offer another advantage: they can be assigned real time units. Because we use the capture times as prior information, this places the inferred latent time on the same scale as the capture times. That is, with appropriate normalization, we can report the latent times in units of days or hours. Consequently, the rate parameters can also be assigned units, such as transcripts per minute. To explore this, we converted the latent time values into units of minutes and normalized the transcription rate parameters to the absolute number of transcripts in a typical mammalian cell. This shows that the transcription rates are roughly on the order of 1-10 transcripts per minute for most genes (Fig. S13), in accordance with previous estimates from single-molecule FISH experiments [11]. The splicing rates are predicted to be slightly lower, on the order of 0.1-1 transcripts per minute. This is consistent with a previous study that estimated the elongation rate in human cells at 3.8 kb/min and found that splicing begins within 10 min [30]. Our estimated degradation rates are similar to the transcription rates, on the order of 1-10 transcripts per minute. Though these estimates should not be taken as definitive, we find it reassuring that our estimated rate parameters are on roughly the correct order of magnitude compared with prior knowledge.

## 3 Discussion

The VeloVAE model uses variational Bayesian inference to estimate cell differentiation progress, cell state, and kinetic rate parameters in a statistically principled fashion. Our approach not only improves gene fitting in many cases, but also resolves some of the key limitations of previous RNA velocity methods. The expressiveness of neural networks makes it possible to adapt the model to various types of gene expression kinetics in biological differentiation systems of varying complexity. Moreover, this model expressiveness does not come at the cost of interpretability; we still learn a set of rate parameters with direct biophysical interpretations. Another key advantage of VeloVAE is that it can perform inference using only the unspliced and spliced mRNA counts from a set of cells, but is also able to incorporate additional prior information such as cell capture times when available. Additionally, the use of mini-batch stochastic gradient descent allows the method to process large datasets without loading all cells into memory at once.

These advantages of VeloVAE make the RNA velocity results much more useful and interpretable in several ways. First, the latent times inferred by the model are sufficiently accurate that they can be used to order cells according to differentiation progress–with even higher accuracy than pseudotime inference in some cases. Second, the ability to use cell capture times as a prior distribution places the inferred cell times on a time scale with known units (e.g., hours or days). Knowing the time units also gives the kinetic rate parameters a more direct interpretation. Furthermore, the fitted and extrapolated values are now qualitatively accurate for individual genes–in contrast to results from previous RNA velocity methods, where the individual gene fits are often poor even when the global latent time and stream plots are reasonable. This opens up many new applications in which knowing the direction and rate of change for an individual gene is important, rather than simply a qualitative, global visualization. For example, one could now perform direct comparison of the times at which different genes are transcribed, revealing the relative order in which genes are activated. Additionally, we showed that the latent time results and future cell state predictions are sufficiently accurate that they can be used to infer transition graphs among cell types in many cases. Finally, our branching ODE model provides important insights into how transcription, splicing, and degradation rates change during cell type bifurcations, which holds promise for understanding the factors regulating cell fate decisions.

## 4 Code and Data Availability

Our code is available at https://github.com/welch-lab/VeloVAE. All datasets analyzed in the paper are previously published and freely available.

## 6 Acknowledgements

This work is funded by NIH grant R01HG010883 to JDW. We would like to thank Chen Li for helpful discussions and Jian Shu, Koseki Kobayashi-Kirschvink and Chengxiang Qiu for help with preprocessing the iPSC reprogramming dataset.

## 7 Methods

### 7.1 Problem Setup

We are interested in the following problem that arises in the context of modeling cellular gene expression changes. Each sample (cell), indexed by *i*, is represented by a vector *X_i_*(*t*) ∈ ℝ^*d*^ parametrized by time *t*. The trajectory *X_i_*(*t*) is governed by some differential equation plus random noise. However, for each *i*, only the vector *x_i_*: = *X_i_*(*t_i_*) is observed at some unknown time *t_i_*. Our goal is two-fold: recover the latent time *t_i_* for each sample and predict future states, i.e., *X_i_* (*t*) for *t* > *t_i_*.

The key biochemical insight underlying our approach is that to express a gene, two types of RNA, nascent unspliced and mature spliced RNA, are produced sequentially. Increases in the unspliced count (*u*) for a gene must precede increases in the spliced count (*s*). This simple insight makes it possible to recover the ordering of cells lacking time labels.

We assume that a dynamical system *F*(*t*; ***θ***) generates scRNA count data. Here, ***θ*** is a set of parameters describing the system, such as the transcription, splicing and degradation rates. Our goal is to use observed scRNA data to simultaneously estimate the parameters ***θ*** of *F* and infer the unknown cell times *t*.

#### Definition 1.

*Let u_g_ and s_g_ denote the unspliced and spliced mRNA count of the g-th gene. Let* 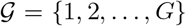 *be a set of genes measured in an scRNA-seq experiment. The **feature vector** of a cell is defined as* **x** = [*u*_1_, *u*_2_, …, *u_G_*, *s*_1_, *s*_2_ …, *s_G_*]^*T*^.

#### Definition 2.

*The **kinetic equation** of gene g is defined as a system of ordinary differential equations relating changes in u and s over time. If there exists a solution F*(*t; **θ***) *to the initial value problem with u*(0) = *u*_0_, *s*(0) = *s*_0_, *we call this solution the **kinetic function** for g.*

#### Definition 3.

*Given a kinetic function u*(*t*) *and s*(*t*) *of a gene, the **RNA velocity** of the gene is defined as* 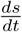.

### 7.2 Modeling Gene Expression Kinetics

In previous work [19], the kinetic equation is modeled by a system of two linear ODEs:

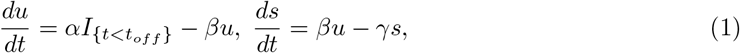

where *I*_{·}_ is an indicator function for the condition in brackets. The model parameters *α, β* and *γ* correspond to the RNA transcription, splicing and degradation rates, respectively. The model assumes that two discrete phases can occur in the gene expression process: (1) induction, when new unspliced RNA molecules are being transcribed (2) repression, when the transcription process stops and no new unspliced molecules are made. The induction phase is assumed to start at *t_on_* = 0 and the transition from induction to repression occurs at time *t_off_*. Given an initial condition *u*(0) = *u*_0_, *s*(0) = *s*_0_, the analytical solution to the ODE is

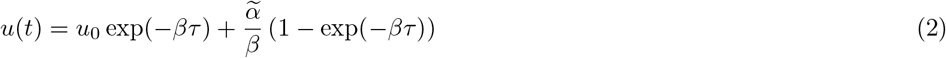

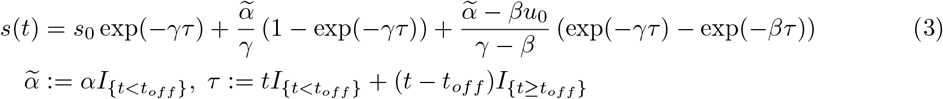

### 7.3 Variational Mixture of ODE Model

#### ODE Formulation

We adopt an ODE formulation similar to (1), except that the transcription rate for each gene is not a single constant *α* anymore. Instead, we assume that the kinetic equation is a continuous mixture of ODEs with transcription rate parameters 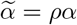. The relative transcription rate *ρ* ∈ [0,1] is a function of latent cell state c, and thus may be slightly different in each cell. The new kinetic equation is:

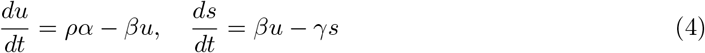

Note that there are no longer discrete induction and repression phases. This can be viewed as a generalization of (1), since *ρ* =1 and *ρ* = 0 correspond to the discrete induction and repression phases, respectively, used in the simpler formulation. Because *ρ* is constant with respect to time, we can still solve the kinetic equation analytically to obtain a closed form for the kinetic function *F*(*t; **θ***) in terms of *α, β, γ*, and *ρ*. The solution is the same as (2) and (3) except that 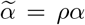. Note also that for each gene, *α, β*, and *γ* are still shared across cells. This model can now capture continuous transcription changes such as those in a bifurcating developmental process.

#### Generative Process

The generative process for the variational mixture of ODE model is as follows:

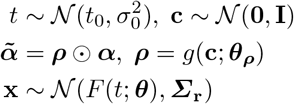

Here, *g*(·) is a neural network with parameters ***θ_ρ_***, ⊙ is the elementwise product, *F* is the kinetic function of all genes, and ***Σ***_**r**_ is a diagonal covariance matrix. This generative process relies on a function *g* mapping latent cell states **c** to relative transcription rates ***ρ***. Intuitively, the cell states can model continuous and bifurcating developmental paths, allowing the entire set of cells to be described as a family of ODEs whose parameters vary smoothly over the cell state manifold. Even though the mapping function *g* is unknown and the posterior of **c** and *t* is intractable, we can apply variational inference to fit a variational mixture of ODEs.

#### Parameter Inference and Neural Network Architecture

We train VeloVAE using the standard mini-batch stochastic gradient descent. The training objective function is the evidence lower bound (ELBO):

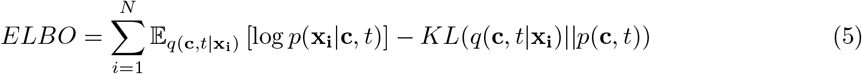

We assume the prior *p*(**c**,*t*) = *p*(**c**)*p*(*t*) is a multi-variate Gaussian distribution where *p*(*c*) is isotropic Gaussian and 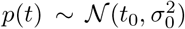. However, the model can take an informative time prior 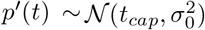 if a cell-wise capture time *t_cap_* is available. In this case, *σ*_0_ is proportional to the time interval between two adjacent capture time points.

The encoder of VeloVAE is a multi-layer perceptron (MLP) with 2 hidden layers and 4 output layers representing the variational posterior mean and standard deviation of **c** and *t*. We use an MLP that is the mirror image of *h* (two layers with 250 and 500 neurons, respectively) to learn the mapping g from **c** to ***ρ***.

Note that we use stochastic gradient descent (SGD) to estimate the ODE rate parameters, unlike scVelo, which used the Nelder-Mead simplex algorithm [24]. The use of SGD instead of Nelder-Mead allows us to use a unified optimization strategy for neural network and ODE parameters, as well as to avoid using all cells at each iteration by loading individual minibatches.

#### Initial Conditions

Because each cell potentially has different ODE parameters, determining the initial conditions is more complex than in the scVelo model. Thus, instead of making the initial conditions trainable parameters, we simply train the model with *u*_0_ = *s*_0_ = 0 in all of our experiments. This still yields excellent data reconstruction and latent time inference. However, the initial conditions are important for accurately predicting the future state of each cell. To improve the accuracy of future state prediction, we first train the VeloVAE to convergence using *u*_0_ = *s*_0_ = 0 so that latent times and cell states are accurate, then determine the initial conditions for a cell at time *t* by simply averaging the (*u, s*) values observed in an immediately preceding time interval [*t* – *δ*_1_, *t* – *δ*_2_]. We then fine-tune the ODE parameters using these updated initial conditions, keeping latent time and cell state fixed.

#### Estimating ODE Parameter Uncertainty

The VAE model can be extended [16] to account for uncertainty in the parameters of the generative model, which in our case are the kinetic rate parameters of the ODE system. Let ***θ*** be the set of all ODE parameters. We assume that some prior distribution *p*_**λ**_(***θ***) generates the parameters. Similar to the (intractable) problem of inferring the latent cell time and state variables, we can use a variational approximation of the posterior *q*_*φ*_(***θ***). The marginal likelihood of the input features can be bounded by

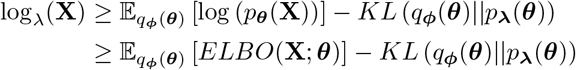

For the prior *p*_**λ**_(***θ***), we choose a factorized log-normal distribution, i.e. each rate parameter *α, β* or *γ* is a random variable drawn from a log-normal distribution. By default, the logarithm of rate parameters has a mean of zero and a standard deviation of 1 for *α* and 0.5 for *β* and *γ*. The posterior mean is initialized using the same method described in the previous paragraph. Again, we can apply the reparameterization trick to take samples of ***θ*** and estimate the expectation of ELBO with a sample mean. The KL divergence between *q*_***φ***_(***θ***) and *p*_**λ**_(***θ***) can be viewed as a regularization of the ODE parameters.

This statistical approach also has a nice intuition in our case: it prevents the “vanishing velocity” problem. If we don’t regularize (*α, β, γ*), optimizing the evidence lower bound may lead to large rate parameters, so that 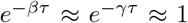. This in turn causes the velocity to vanish so that each cell will be approximately at the steady state, with 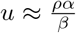 and 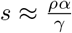. Because *ρ* is given by a neural network *g*, we can achieve good reconstruction if *g* is expressive enough, but the velocity will be close to zero. This issue is resolved by placing a prior distribution on the rates and including them in the variational approximation. This has the effect of regularizing the posterior distribution of the rates toward their prior, avoiding the vanishing velocity problem.

#### Interpreting Rate Parameters

Although the model accepts any notion of time and rates, the units can be converted to match the actual units of cell developmental time, usually in days. Suppose the cell time from the model ranges from *t*_0_ to *t*_1_, in a hypothetical time unit, and we have prior knowledge about the entire duration of *m* days. From the mathematical property of our ODE system, we know that scaling time by *k* and rate parameters (*α,β,γ*) by 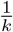 results in the same u and s. Thus, we can perform unit analysis to convert the rates into units of minutes as follows:

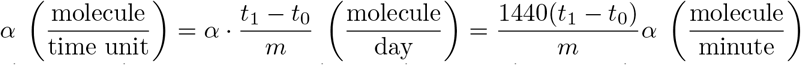

Similarly, 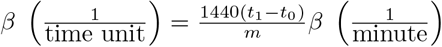 and 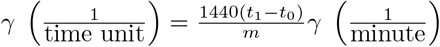

We can then scale the rate parameters to account for the fact that scRNA-seq captures only a fraction of the transcripts in a cell. A typical mammalian cell contains about 360000 mRNAs^4^. Therefore, we can scale *ρ* by 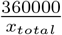, where *x_total_* is the median total mRNA count number. We analyzed rate parameters learned from our model, converted the units and computed the histograms (Supplementary Figure S13). For *α*, we analyzed the peak transcription rate by considering the case of highest transcription (*ρ* = 1). For *β* and *γ*, we multiplied them by *u_top_* and *s_top_* respectively to obtain the same units as *α*. Here, *u_top_* and *s_top_* are the 95-percentile *u* and *s* values, representing cells with high expression levels.

### 7.4 Data Preprocessing

We loaded all datasets from AnnData (h5ad) format. We first used scanpy [38] to select highly variable genes, normalize and scale the gene expression counts. Next, we followed the scVelo preprocessing pipeline by performing principal component analysis on the normalized and scaled expression data, then smoothing the unspliced and spliced expression levels among k-nearest neighbors identified from the principal components.

### 7.5 Parameter Initialization

The neural network weights are randomly initialized with Xavier uniform distributions. The only exception is the output layer of the decoder network, which is initialized with Xavier normal distributions, to prevent gradient vanishing of the sigmoid activation. VeloVAE applies the same initialization method as scVelo by applying the steady-state assumption and the dynamical model. Next, a global cell time is estimated as the median of locally initialized time across all genes.

If capture times are available, VeloVAE initializes a unique cell time by directly sampling from a distribution centered at the real time. Whether the initial global time is estimated by the steadystate model or directly sampling from capture time, rate parameters are always re-estimated based on the dynamical model. We still make the assumption that *β* = 1. First, switch-off time is estimated as the average global cell time of the steady-state cells. Next, we estimate *α* and the switch-on time *t_on_* by solving a set of equations using two cells with their initialized cell time (*u*_1_,*t*_1_), (*u*_2_,*t*_2_):

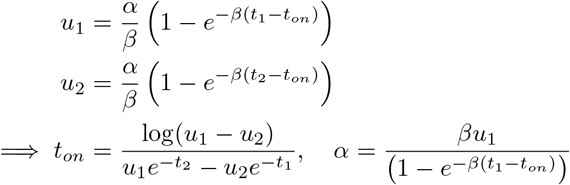

Note that *u*_1_ and *u*_2_ are chosen as the sample average around the median and top quantile of u values to promote robustness against noise. Finally, *γ* is estimated by 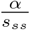 where *s_ss_* is the estimated steady-state *s* value.

### 7.6 Branching ODE

#### Model Description

A variational mixture of ODEs provides sufficient flexibility to account for complex kinetics, but only describes cell-wise kinetics. In order to distill qualitative knowledge about gene expression kinetics and reveal cell-type relations, we propose a new ODE model called branching ODE.

The fundamental assumptions we make about branching ODE include:

1. Each cell belongs to exactly one of the cell types *y*_1_, …, *y_k_*.
2. At least one of the cell types is a stem cell type. Each non-stem-cell type is a descendant of exactly one other cell type. (Note that we could relax this assumption to allow multiple progenitors, but we choose not to purely for simplicity.) Each cell type has an initial time *t*_0_ when it emerges in the differentiation process.
3. Cells of the same type, *y*, share identical transcription, splicing and degradation rates (*α_y_, β_y_, γ_y_*).

With these assumptions, we can summarize differentiation with directed cell type transition graph *G* = (*V* = {*v*_1_,…, *v_k_*}, *E*), called a transition graph. Each cell type corresponds to a vertex in *G*, and each edge (*u, v*) represents the relation of *u* differentiating into *v*. By our second assumption, *G* is composed of one or multiple trees. The kinetic functions retain the same analytical form, except for type-specific rate parameters and initial conditions. Furthermore, the equations are defined recursively because the initial condition of any non-stem-cell type depends on its progenitor cell type.

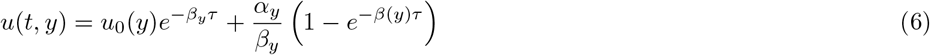

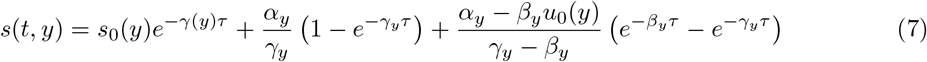

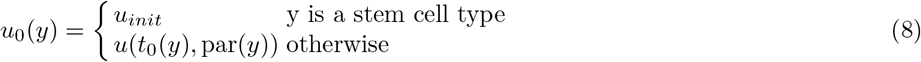

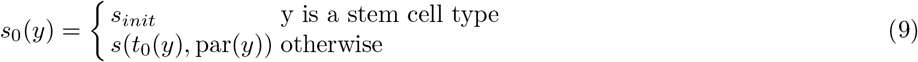

Here, par(·) refers to the parent vertex in the transition graph. Note that *τ* is redefined as *τ*: = *t* – *t*_0_(*y*) for each cell type *y*, i.e. *τ* is the time duration starting from the initial time of each cell type.

#### Inferring the Transition Graph

A challenging problem for applying branching ODE is the absence of the transition graph. In many cases, we do have some prior knowledge of the transition graph, but it is more desirable to simply infer the graph directly from the data when possible. Therefore, we apply a computational method to infer the transition graph based on cell time and states. Because the transition graph is a collection of arborescences, i.e. rooted trees in a directed graph, we can solve the problem with simple graph algorithms.

First, we partition the cells into distinct cluster(s). In particular, we perform Leiden clustering with a low resolution on the UMAP coordinates of the data.

Next, we build a complete subgraph in each partition. Let 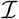 be the set of all cells and 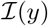 be the set of all cells of type *y* in a partition. For each cell 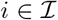 with time *t_i_*, we take a time window [*t_i_* – *δ*_1_, *t_i_* – *δ*_2_] and find k nearest neighbors based on cell state **c**. Denote 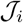 as the set of neighbors of *i*. Then, for any two cell types *y* and *z*, the empirical transition probability from *y* to *z* is defined as

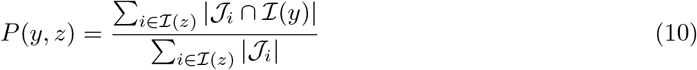

In other words, we group cells into the cell types and count the number of transitions from any cell type to any other type.

Finally, we apply Edmond’s algorithm [9] to find the maximum spanning arborescence in each partition. The earliest cell type is choosen to be the root. The algorithm starts by picking the parent *y* of each non-root vertex *z* greedily, i.e. *y* = argmax_*v*_ *P*(*v, z*). Next, it checks loops, collapses loops into super-vertices and is recursively applied to the new graph until no loop exists. Finally, it breaks the loops after each level of recursion.

#### Training the Branching ODE

We assume that the cell time has already been inferred from VeloVAE. Thus, the branching ODE is a regression model with an analytical form shown in equation (6), (7). To train the model, we find the cell-type-specific rate parameters ***θ*** that maximize the Gaussian likelihood, which is equivalent to minimizing a Mahalanobis distance:

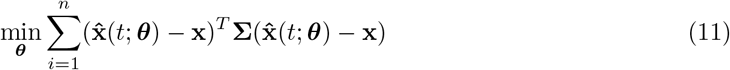

Here, 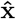 is predicted expression level. We train using mini-batch stochastic gradient descent with the ADAM optimizer.

---

**Fig. S1.**
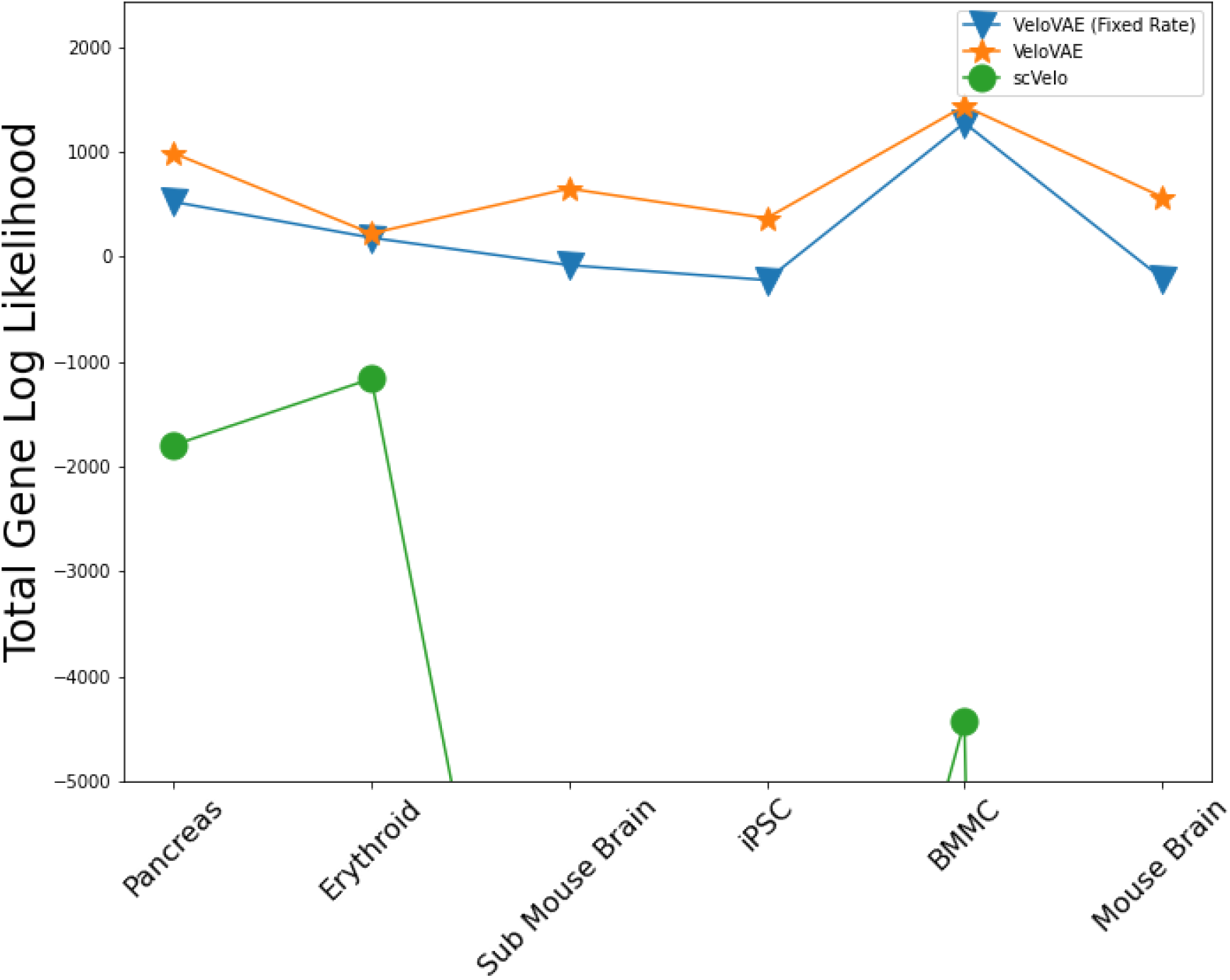
Comparison of fitted gene log likelihood. We compared VeloVAE with fixed rate (only time in the latent space), VeloVAE and scVelo. The gene total likelihood assumes Gaussian likelihood and is computed by taking the mean of the sum of gene log likelihood. The y-axis is truncated to better show the difference between VeloVAE (fixed rate) and VeloVAE. Note that we only compared the genes which scVelo fitted with a positive likelihood, as there are genes scVelo didn’t fit or fitted with zero likelihood.

**Fig. S2.**
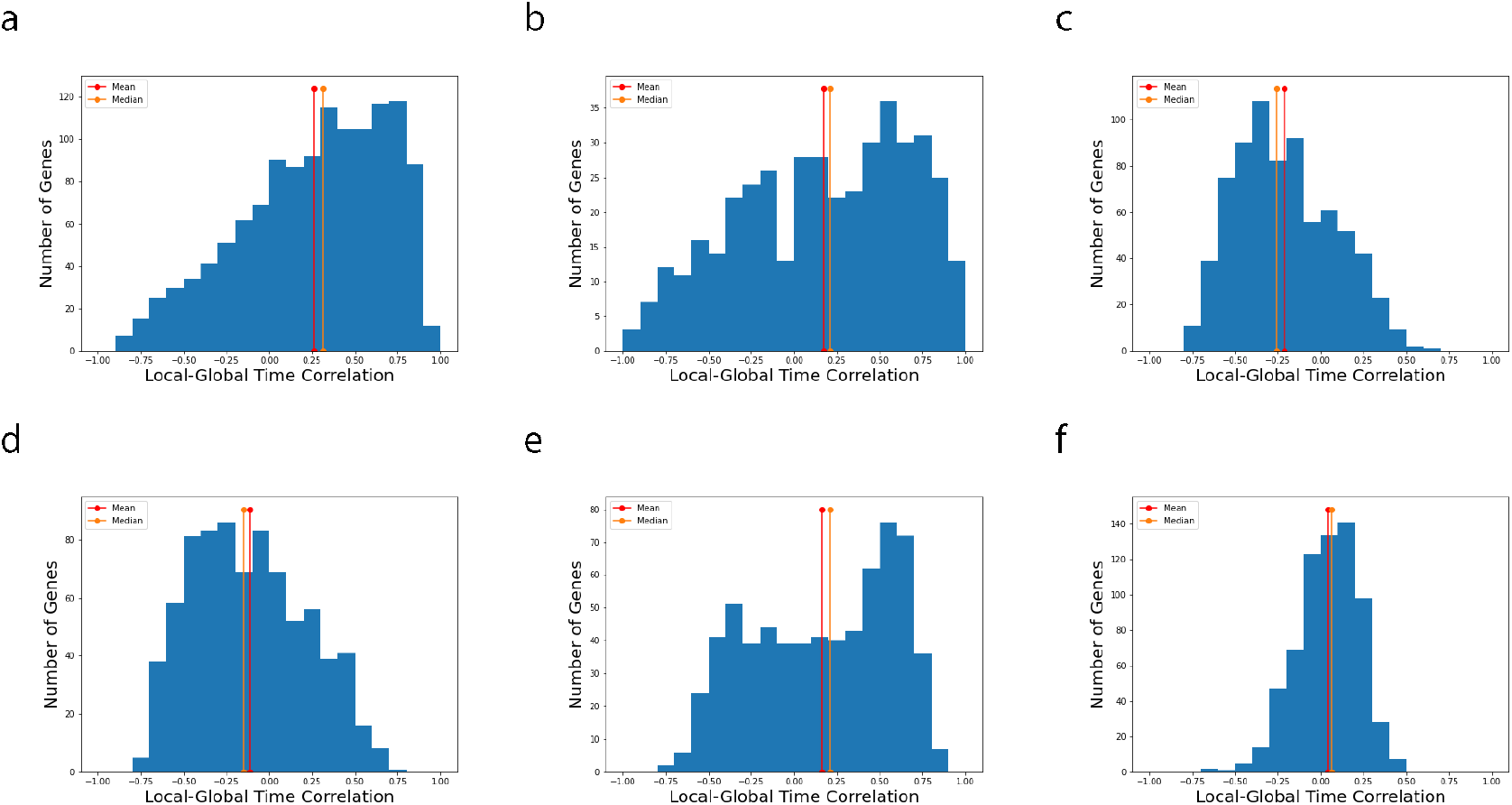
Histogram of scVelo time correlation. We computed the the Spearman correlation between scVelo locally fitted (gene-specific) time and scVelo global latent time for six datasets. This gives a correlation value for each fitted gene. The figures shows a histogram of these correlations for pancreas **(a)**, erythroid **(b)**, subsampled mouse brain **(c)**, iPSC **(d)**, human bone marrow **(e)** and full mouse brain **(f)**.

**Fig. S3.**
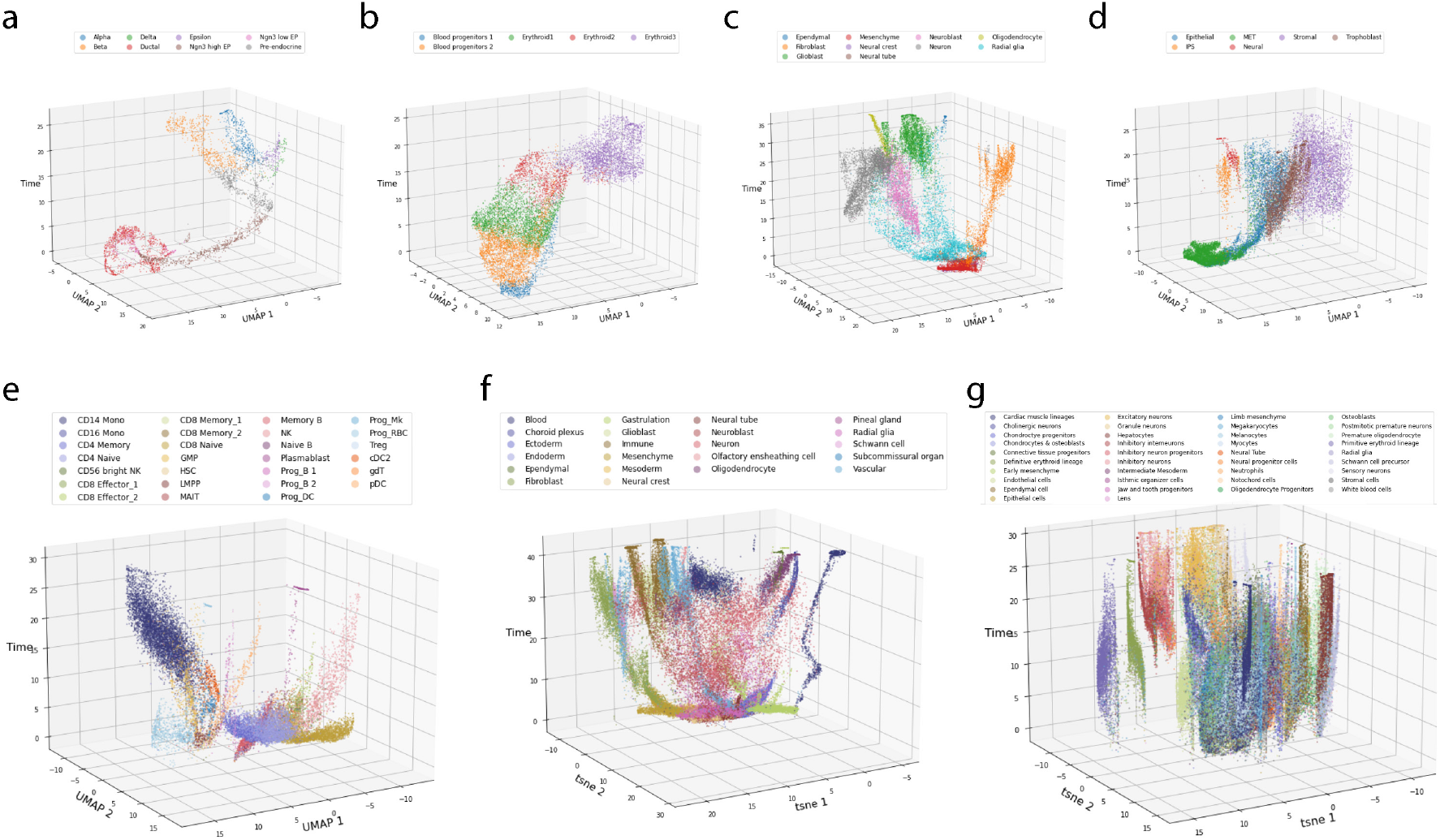
Cell state-time plot. 2D UMAP embeddings of the latent cell state **c** plotted versus inferred latent time. Each plot has the 2D UMAP embedding as the x-y plane and cell time as the z axis. The plots can be interpreted as time evolution of the cell state space. The seven panels correspond to pancreas **(a)**, erythroid **(b)**, subsampled mouse brain **(c)**, iPSC **(d)**, human bone marrow **(e)**, full mouse brain **(f)** and mouse embryo **(g)**.

**Fig. S4.**
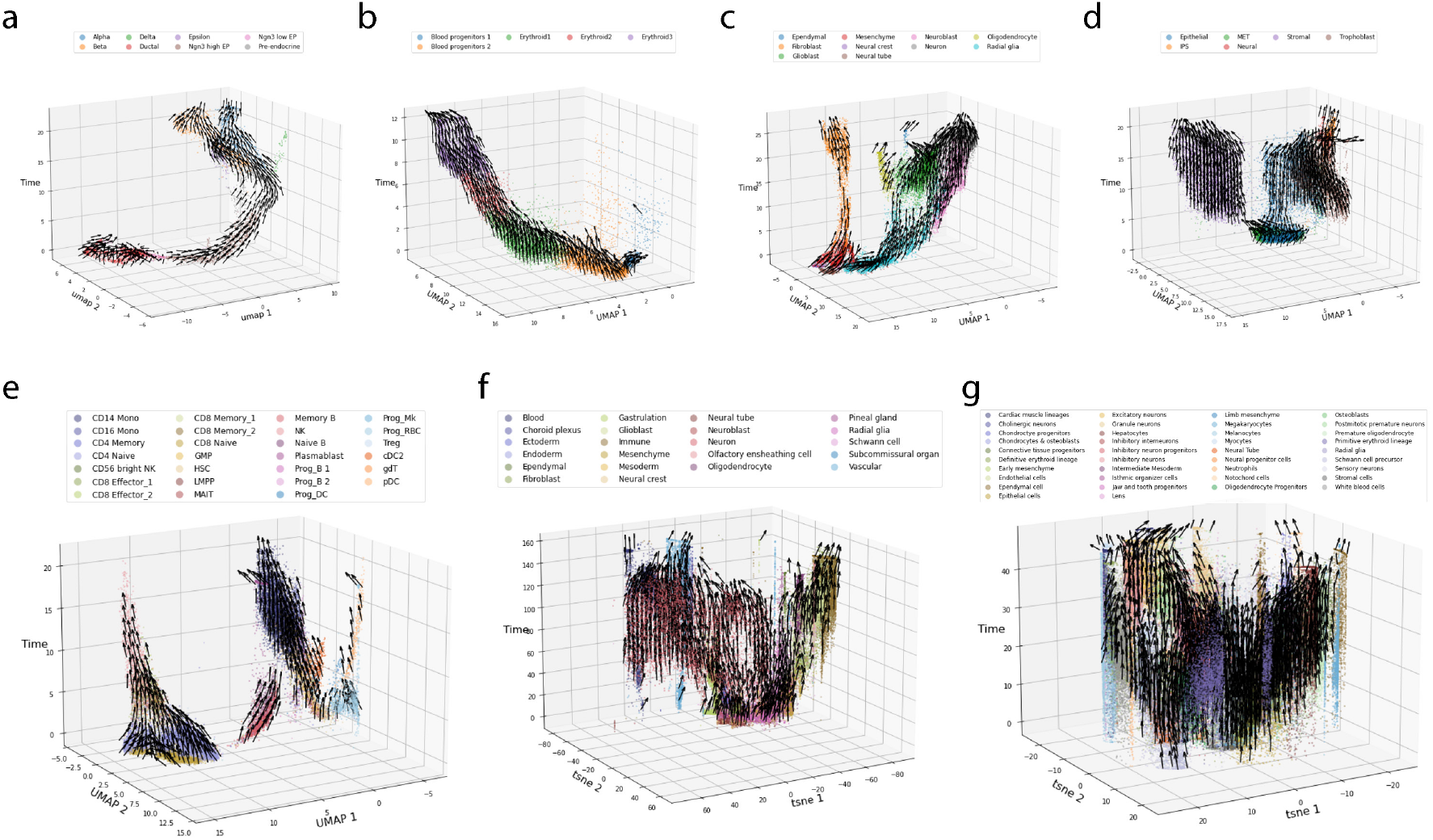
3D velocity plot. 3D plots computed as described in Fig. S3, with velocity vectors added. The x-y plane is either UMAP (a-e) or t-SNE (f-g) coordinates. The 3D embedding is computed by averaging displacement vectors towards k neareast neighbors in the immediate future, i.e. arrows point upwards along the z axis. The seven panels correspond to pancreas **(a)**, erythroid **(b)**, subsampled mouse brain**(c)**, iPSC **(d)**, human bone marrow **(e)**, full mouse brain **(f)** and mouse embryo **(g)**.

**Fig. S5.**
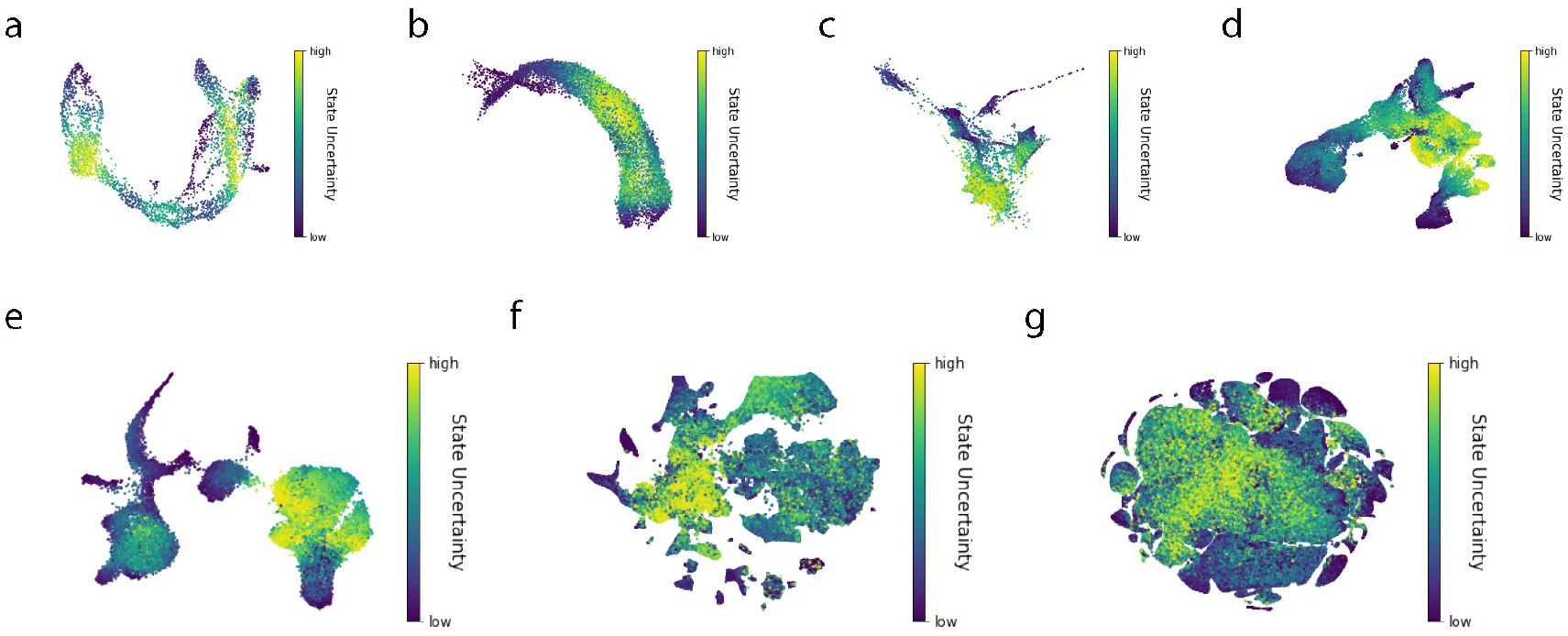
Inferred cell state uncertainty. Each panel shows a UMAP or t-SNE plot colored by the multi-variate coefficient of variation of the cell state. Cell state is a continuous identification of cell type, so conceptually it should have high uncertainty in progenitor cell types, as cell fate is undetermined. Our results verified this intuition as well as the capability of learning meaningful representations using VeloVAE. The 7 panels correspond to pancreas **(a)**, erythroid **(b)**, subsampled mouse brain **(c)**, iPSC **(d)**, human bone marrow **(e)**, full mouse brain **(f)** and mouse embryo **(g)**.

**Fig. S6.**
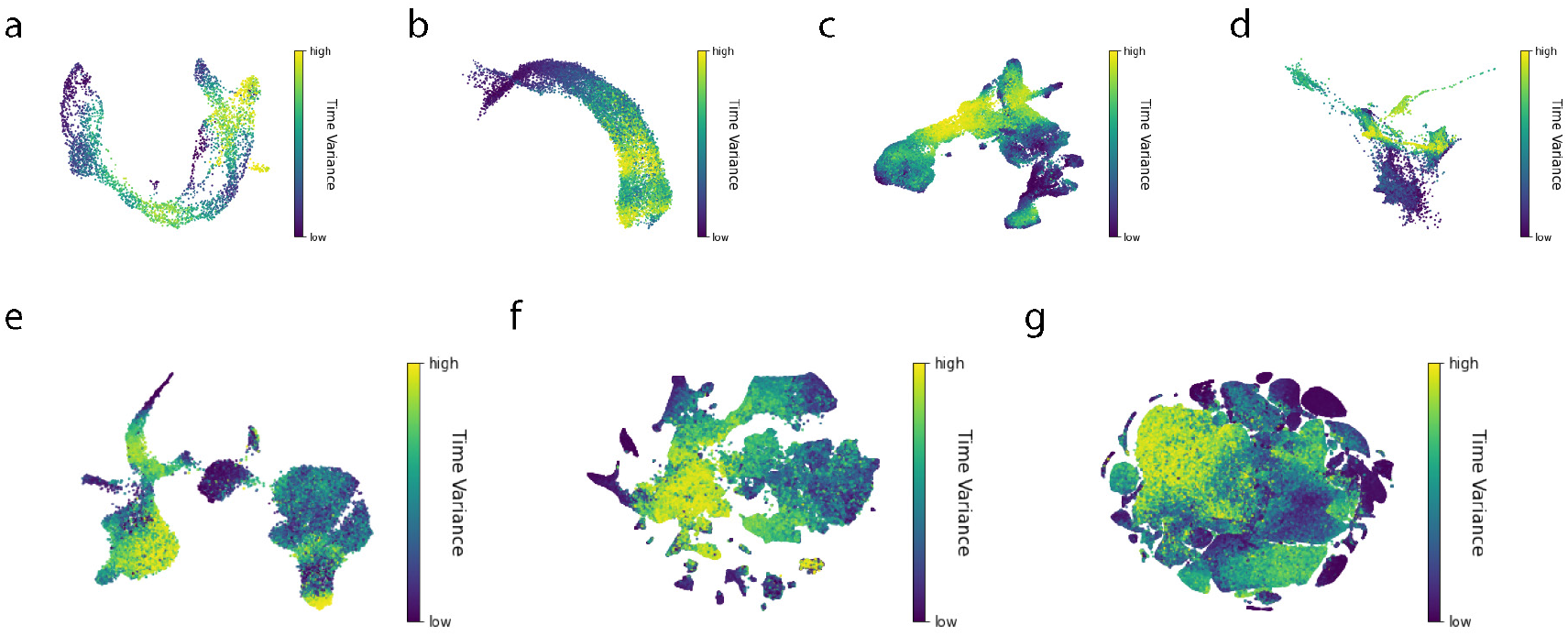
Time variance in test datasets. Each panel shows a UMAP or t-SNE plot colored by the coefficient of variation of the inferred cell time. The 7 panels correspond to pancreas **(a)**, erythroid **(b)**, subsampled mouse brain**(c)**, iPSC **(d)**, human bone marrow **(e)**, full mouse brain **(f)** and mouse embryo **(g)**.

**Fig. S7.**
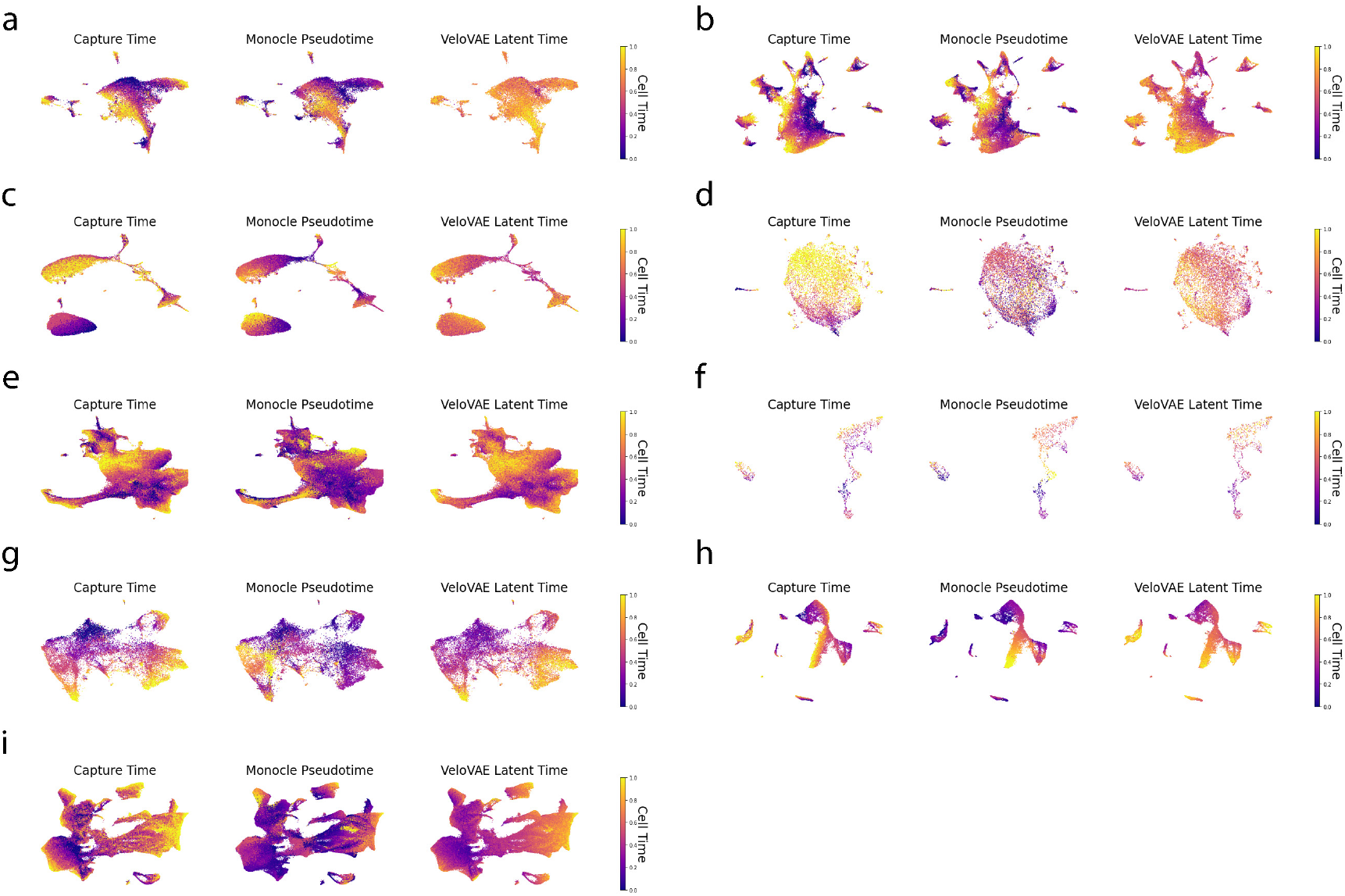
Latent time of major trajectories from the mouse embryo. We compare the capture time, monocle 3 pseudotime and VeloVAE latent time by showing the UMAP plots colored by latent time of ten major trajectories in the embryo dataset: endothelial **(a)**, epithelial **(b)**, haematopoiesis **(c)**, hepatic **(d)**, mesenchymal **(e)**, melanocyte **(f)**, PNS glia **(g)**, PNS neuron **(h)** and neural tube and notochord **(i)** trajectories.

**Fig. S8.**
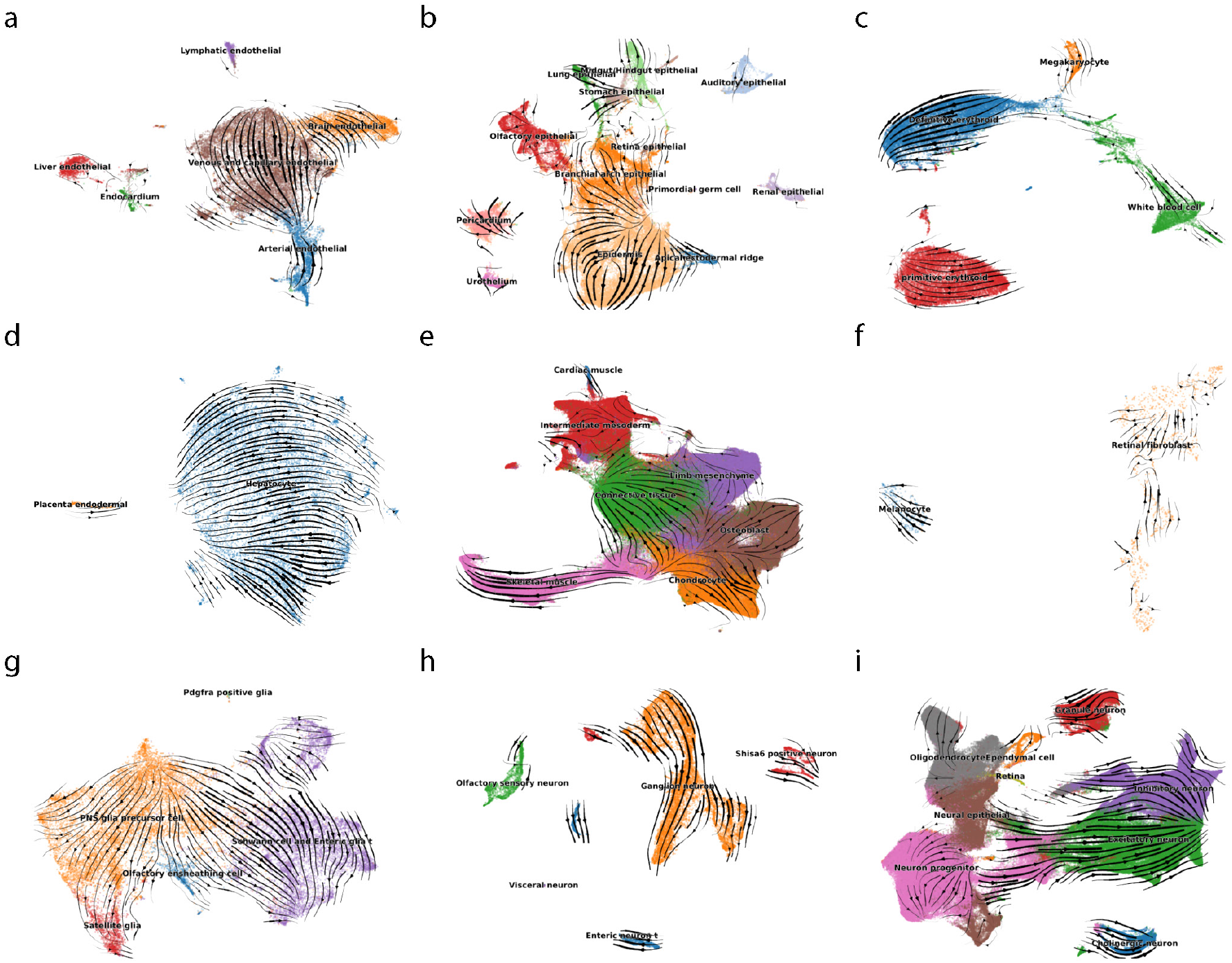
Velocity stream plots for major trajectories from the mouse embryo. We plotted the velocity embedding on UMAP coordinates of ten major trajectories in the embryo dataset: endothelial **(a)**, epithelial **(b)**, haematopoiesis **(c)**, hepatic **(d)**, mesenchymal **(e)**, melanocyte **(f)**, PNS glia**(g)**, PNS neuron **(h)** and neural tube and notochord **(i)** trajectories.

**Fig. S9.**
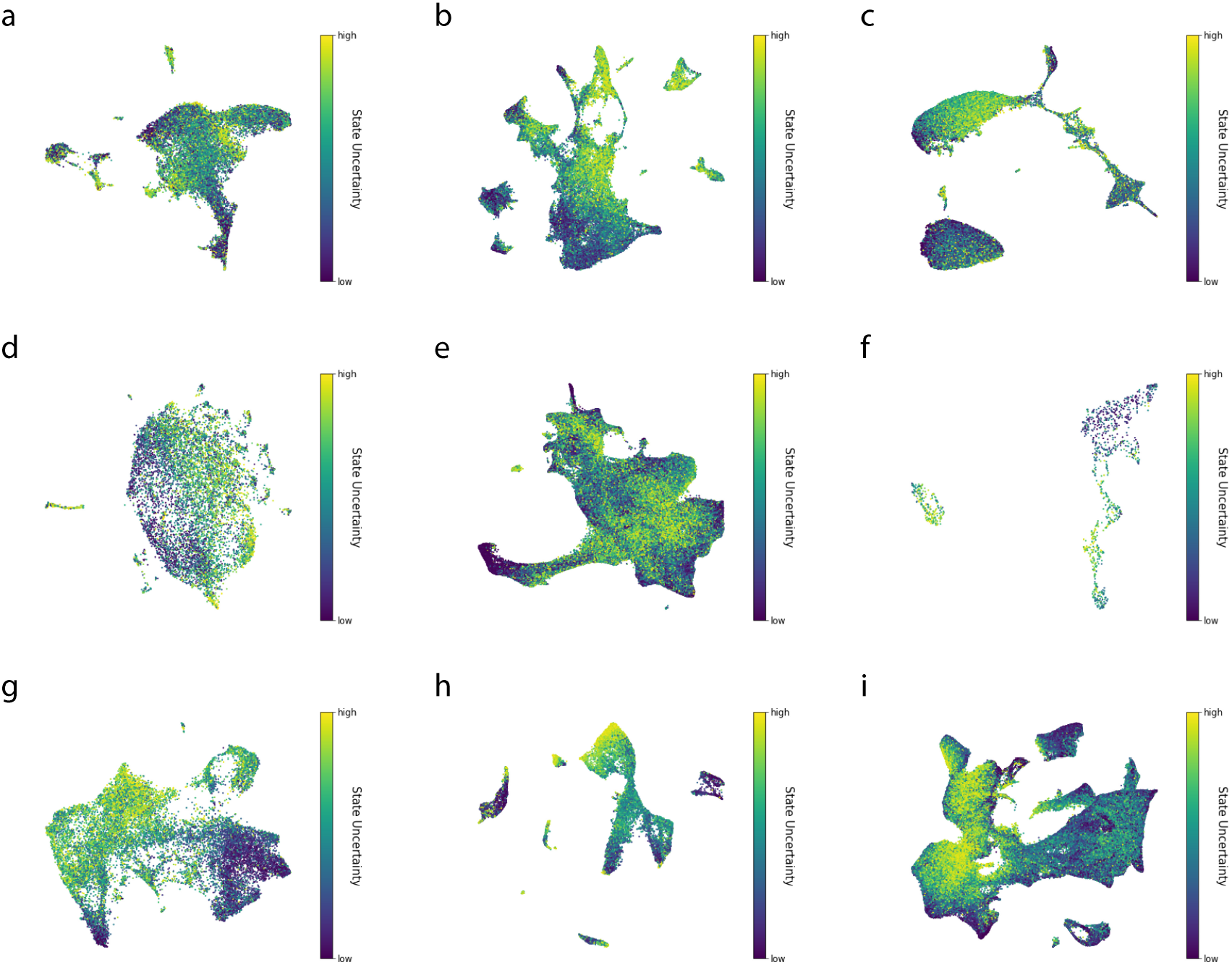
Cell state uncertainty in major mouse embryo trajectories. Each panel shows a UMAP plot colored by the multi-variate coefficient of variation of the cell state. The 10 panels correspond to endothelial **(a)**, epithelial **(b)**, haematopoiesis **(c)**, hepatic **(d)**, mesenchymal **(e)**, melanocyte **(f)**, PNS glia **(g)**, PNS neuron **(h)** and neural tube and notochord**(i)** trajectories in the mouse embryo dataset.

**Fig. S10.**
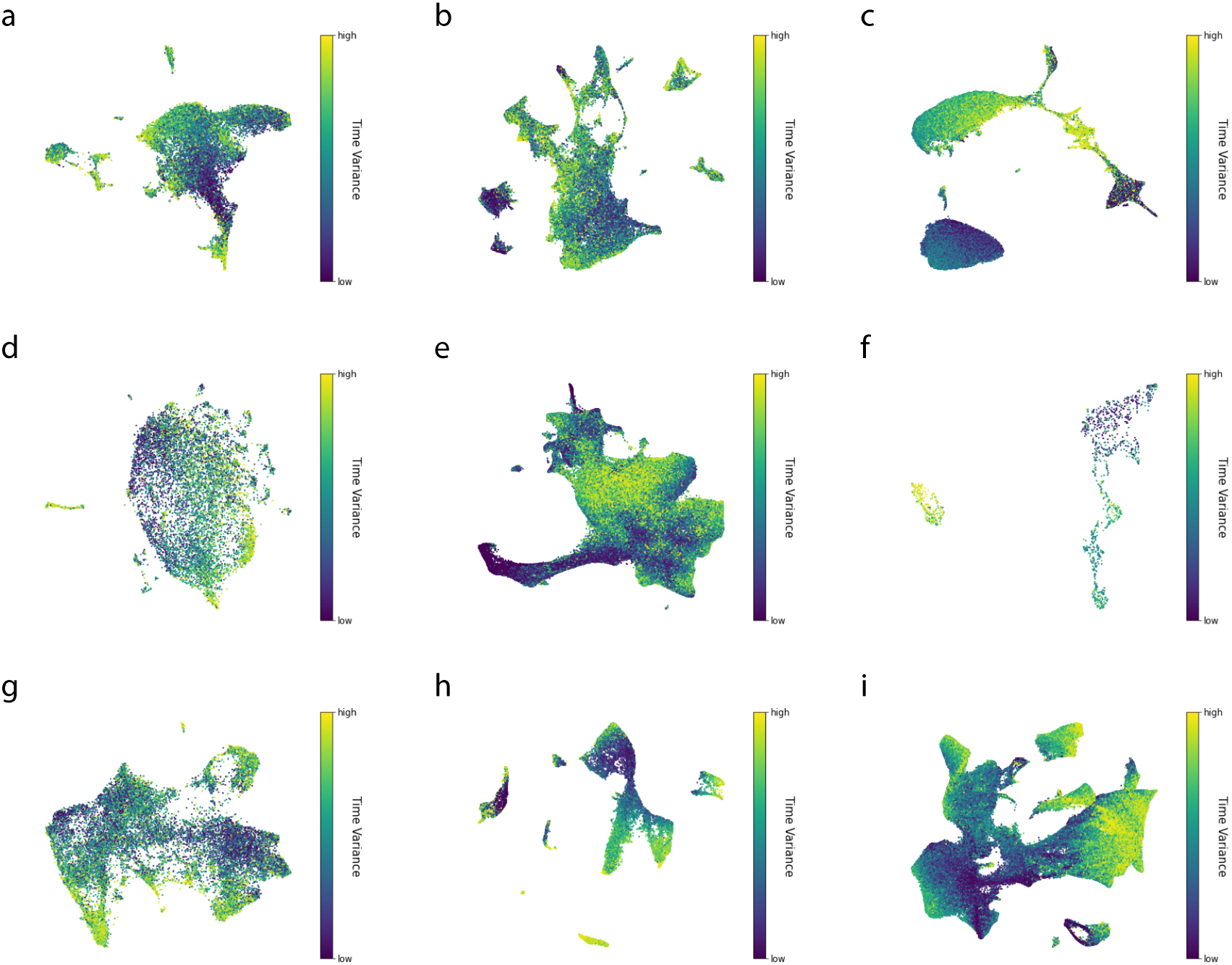
Cell time variance in major mouse embryo trajectories. Each panel shows a UMAP plot colored by the coefficient of variation of the cell time. The 10 panels correspond to endothelial **(a)**, epithelial **(b)**, haematopoiesis **(c)**, hepatic **(d)**, mesenchymal **(e)**, melanocyte **(f)**, PNS glia **(g)**, PNS neuron **(h)** and neural tube and notochord **(i)** trajectories in the mouse embryo dataset.

**Fig. S11.**
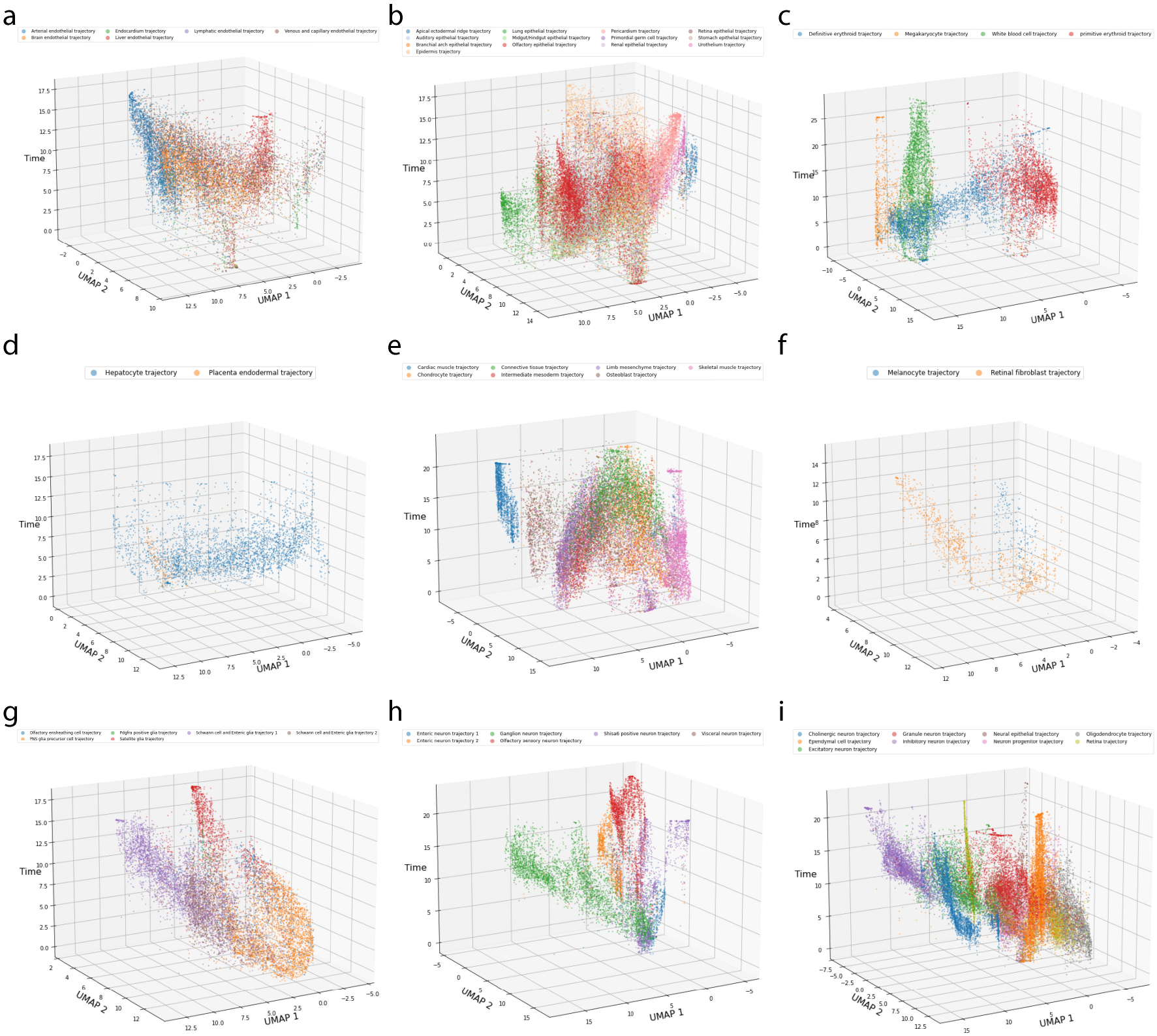
Cell state-time plot. We computed 2D UMAP embeddings of the latent cell state **c** and plot it versus inferred latent time. Following the setting of the original work, we used cosine distance and 15 neighbors. The 10 panels correspond to endothelial **(a)**, epithelial **(b)**, haematopoiesis **(c)**, hepatic **(d)**, mesenchymal **(e)**, melanocyte **(f)**, PNS glia**(g)**, PNS neuron **(h)** and neural tube and notochord **(i)** trajectories in the mouse embryo dataset.

**Fig. S12.**
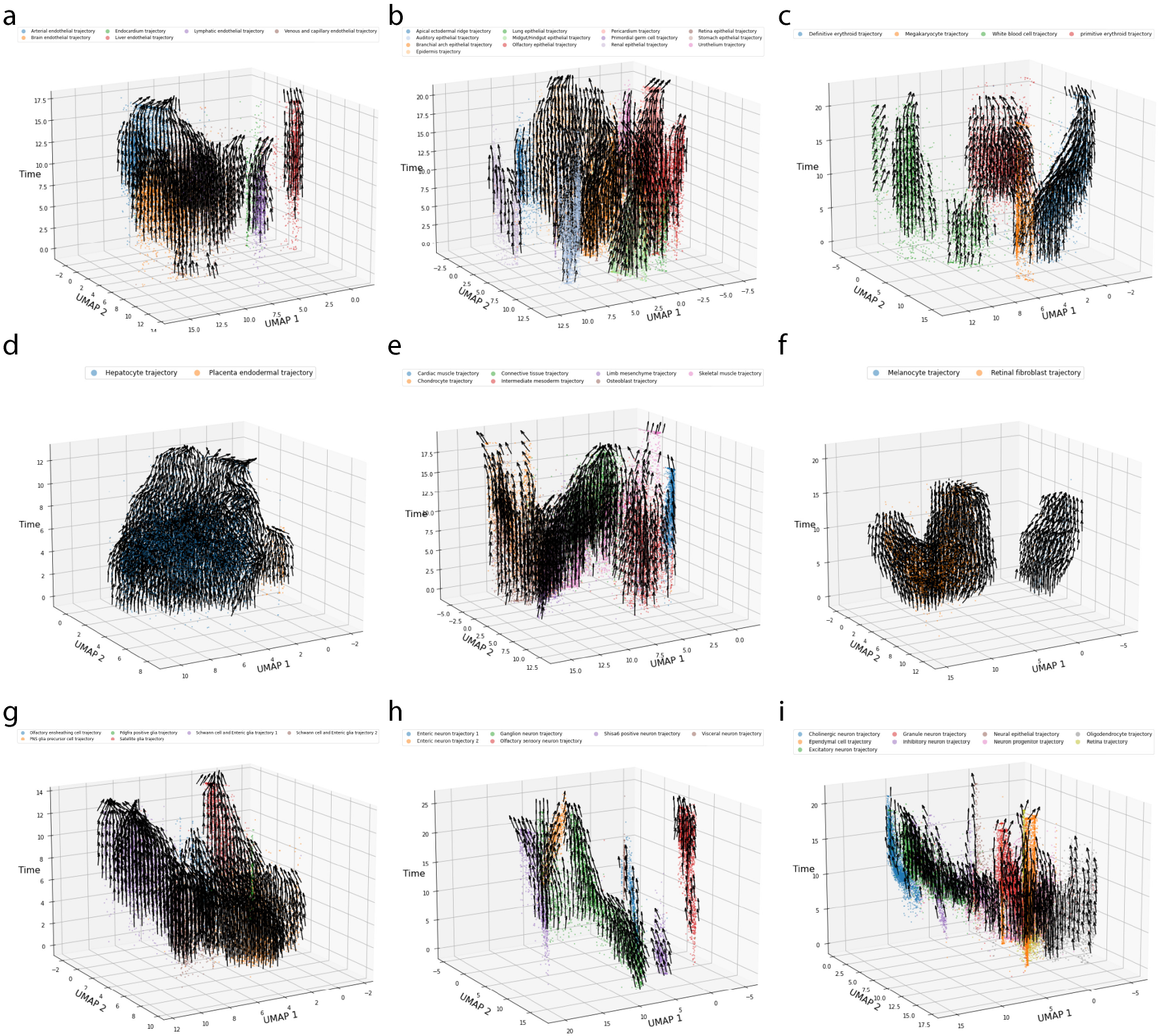
3D velocity plot. 3D velocity plot computed as described in Fig. S4. The 10 panels correspond to endothelial **(a)**, epithelial **(b)**, haematopoiesis **(c)**, hepatic **(d)**, mesenchymal **(e)**, melanocyte **(f)**, PNS glia **(g)**, PNS neuron **(h)** and neural tube and notochord **(i)** trajectories in the mouse embryo dataset.

**Fig. S13.**
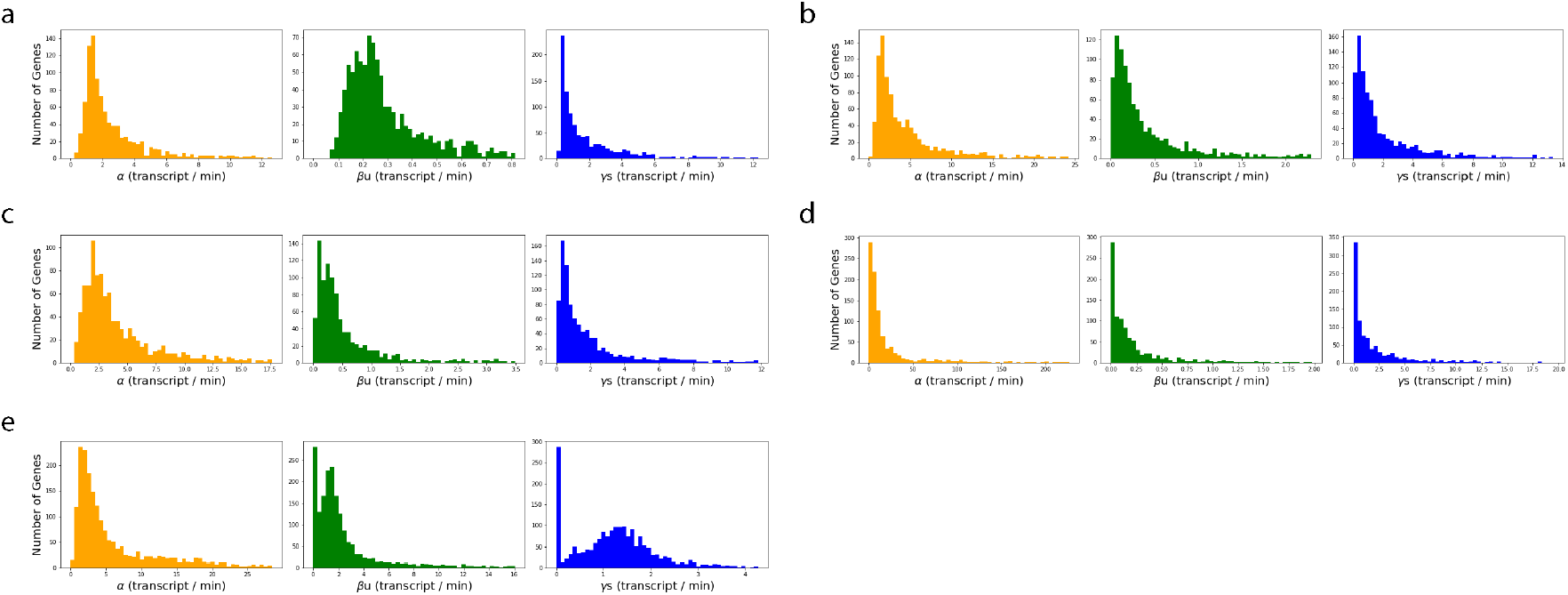
Rate parameter histograms. Histograms of transcription, splicing and degradation rates estimated by VeloVAE. The histogram is computed across genes within each dataset. Transcription rates are reported for *rho* = 1. For splicing and degradation, we report *βu* and *γs* to match the unit of *α* (mRNA / minute). Here, *u* and *s* are chosen to be half of the 95th-percentile count number. The 5 panels correspond to erythroid **(a)**, iPSC **(b)**, subsampled mouse brain **(c)**, full mouse brain **(d)**, and mouse embryo **(i)** datasets.

**Table S1.** Default Hyperparameters of VeloVAE. We trained our model with minimal change to the default hyperparameters. The only changes we made include **(1)** setting early_stop to 9 and train_ton to False for the erythroid dataset **(2)** increasing the batch size to 2048 and n_neighbors to 30 for the mouse embryo dataset.

| Parameter Name | Value | Notes |
| --- | --- | --- |
| $h_1$ | 500 | Width of the first hidden encoder layer/second decoder layer |
| $h_2$ | 250 | Width of the second hidden encoder layer/first decoder layer |
| n_epochs | 1000 | Number of (maximum) training epochs |
| n_epochs_post | 1000 | Number of (maximum) refining epochs |
| batch_size | 128 | Number of samples in a mini-batch |
| learning_rate | $2 \times 10^{-4}$ | Gradient descent learning rate of neural network parameters |
| learning_rate_ode | $5 \times 10^{-4}$ | Gradient descent learning rate of ODE parameters |
| $\lambda$ | $1 \times 10^{-3}$ | L2 regularization coefficient of the encoder network. |
| $\lambda_\rho$ | $1 \times 10^{-3}$ | L2 regularization coefficient of the decoder ρ-network |
| kl_t | 1.0 | time KL divergence coefficient |
| kl_c | 1.0 | cell state KL divergence coefficient |
| test_iter | 2 | number of epochs between two consecutive tests on the validation data |
| n_warmup | 5 | number of initial epochs when only the neural network parameters are updated |
| early_stop | 5 | number of consecutive epochs in the early stopping criteria |
| early_stop_thred | $\# \text{ gene} \times 10^{-3}$ | ELBO change threshold in the early stopping criteria |
| train_test_split | 0.7 | proportion of training samples out of the entire dataset |
| k_alt | 1 | number of iterations before each update of the ODE parameters |
| train_scaling | False | whether to train the scaling parameter of unspliced counts |
| train_std | False | whether to train the standard deviation of Gaussian noise |
| train_ton | True | whether to train the switch-on time of each gene |
| time_overlap | 0.5 | overlap proportion between two informative time prior distributions |
| n_neighbors | 10 | number of neighbors in time-window KNN initial condition estimation |
| dt | (0.03,0.06) | coefficient of the edges of the time window in KNN initial condition estimation |

**Table S2.**
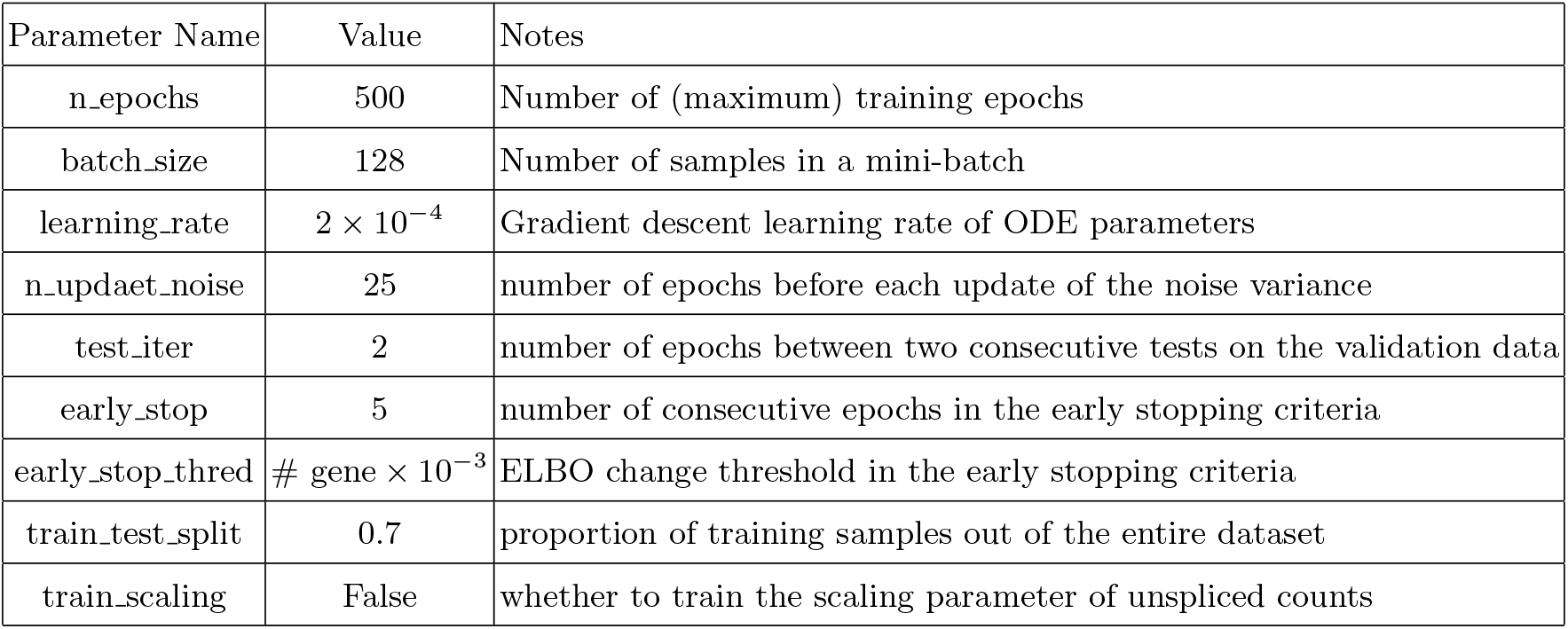
Default Hyperparameters of Branching ODE.

## Footnotes

4 https://www.qiagen.com/us/resources/faq?id=06a192c2-e72d-42e8-9b40-3171e1eb4cb8

